# MacroH2A1 and H3K27me3 target cGAS to facultative heterochromatin to restrain innate immune activation

**DOI:** 10.64898/2026.10.08.757605

**Authors:** Suzie Thenin-Houssier, Cyprien Jahan, Wenjie Xu, Giorgio Papadopoulos, Lucie Bonnet-Madin, Syota Matsumoto, Tomoya Kujirai, Olivier Cuvier, Hitoshi Kurumizaka, Bijan Sobhian, Monsef Benkirane

## Abstract

The DNA sensor cGAS is largely sequestered on chromatin in an inactive form, but how this state is regulated remains unclear. We show that cGAS preferentially binds nucleosomes containing the histone variant macroH2A1 (mH2A1) marked by H3K27me3 and H3K9me3. Cryo-EM reveals that cGAS selectively engages the mH2A1/H2B acidic patch within heterotypic nucleosomes. Depleting mH2A1.1 or blocking H3K27me3 deposition releases cGAS from chromatin and activates the cGAS-STING-IRF3 pathway. Genome-wide profiling reveals two cGAS-bound chromatin compartments: chromosome-arm sites co-occupied by mH2A1.1, methylated CpG islands, and CTCF, where cGAS and mH2A1.1 binding are interdependent; and centromeric regions (14% of peaks), which lack DNA methylation and mH2A1.1 dependence. These findings establish mH2A1 and H3K27me3 as determinants of cGAS chromatin tethering, linking nuclear cGAS regulation to facultative heterochromatin and opening new avenues for modulating cGAS-STING signaling.

---

Major discoveries in recent years have highlighted previously unrecognized physical and functional interactions between two cellular pathways once considered independent: the DNA repair and innate immune pathways^1–7^. On one hand, key DNA repair factors have been demonstrated to act as DNA sensors and/or regulators of DNA sensing^8–14^, while conversely, the major DNA sensor cGAS has been implicated in DNA repair^15–19^. More recently, studies revealed that the majority of cGAS is tightly tethered to chromatin in an enzymatically inactive form^20–22^. Notably, cGAS interaction with the acidic patch of H2A/H2B dimers is approximately 100-fold stronger than its interaction with naked DNA, ensuring stringent nuclear control of cGAS activity^20^. Structural analyses of cGAS bound to in vitro-reconstituted nucleosome core particles (NCPs) revealed that NCPs induce major conformational changes in cGAS, rendering it inactive^23–28^. These changes involve the engagement of critical cGAS residues, normally required for DNA binding, with the H2A/H2B acidic patch, as well as steric hindrance by the NCP that blocks both DNA binding and dimerization sites. These findings challenge our understanding of cellular compartmentalization in the DNA-triggered innate immune response, which traditionally distinguishes self from non-self DNA^16,20,29^. They raise questions about the purpose of cGAS storage in an inactive chromatin-associated form, as well as its regulation and function.

Here, we address these questions by purifying native cGAS-bound nucleosomes from human cells and combining biochemical, structural, and genome-wide approaches. We find that cGAS preferentially associates with heterotypic nucleosomes containing the histone variant macroH2A1 (mH2A1) and marked by the repressive modifications H3K27me3 and H3K9me3. Cryo-EM analysis of a reconstituted cGAS-heterotypic nucleosome complex reveals that cGAS asymmetrically occupies the acidic patch on the mH2A1 side of the particle, providing a structural basis for this selectivity. Depleting mH2A1.1 or pharmacologically inhibiting H3K27me3 deposition releases cGAS from chromatin and triggers cGAS- and STING-dependent activation of the type I interferon response, independently of concurrent DNA damage signaling. Genome-wide profiling further shows that cGAS occupies two epigenetically and mechanistically distinct chromatin compartments: mH2A1.1-dependent sites at chromosome arms, co-occupied by CTCF and DNA methylation, and mH2A1.1-independent centromeric sites marked by constitutive heterochromatin. Together, these findings establish that cGAS chromatin tethering is not a uniform, chromatin-state-independent lock, but is instead directed toward facultative heterochromatin by a specific histone variant and repressive histone mark, thereby coupling nuclear cGAS regulation to heterochromatin identity and genome organization.

## Results

To determine how cGAS is regulated on chromatin and to identify the molecular basis of its nuclear tethering, we sought to purify native cGAS-bound nucleosomes (cGAS-Nuc) directly from human cells. To this end, we developed a chromatin solubilization protocol to purify cGAS-Nuc and their interacting partners, via tandem affinity chromatography^30^. We used a previously well-characterized stable HeLa cGAS knockout (KO) cell line expressing Flag-HA-tagged cGAS (cGAS-FHA) or a control empty vector (EV)^31^. cGAS-FHA was expressed at slightly higher levels than endogenous cGAS (Supplementary Fig. 1a, upper panel), with a comparable subcellular distribution across cytosolic, nuclear soluble, and chromatin fractions (Supplementary Fig. 1a, lower panel; vinculin and Lamin B1 used for fraction purity controls), indicating that epitope tagging does not substantially alter cGAS localization. The chromatin fraction was prepared, subjected to MNase digestion, and extracted with 500 mM NaCl for solubilization (see Methods). Analysis of MNase-digested chromatin showed that DNA fragments in the chromatin extract (ChE) corresponded primarily to mono- and di-nucleosomes (Supplementary Fig. 1b).

Next, ChE were subjected to tandem affinity chromatography using anti-Flag and anti-HA antibodies^30^. Purified cGAS-FHA-Nuc and Flag-HA controls were resolved by SDS-PAGE and silver-stained (Fig. 1a). Associated partners and histone modifications were identified via tandem mass spectrometry (Supplementary Table 1). As this proteomic survey is qualitative rather than quantitative, proteins uniquely present in cGAS-FHA-Nuc eluates with over 40% peptide coverage, were considered candidate interactors and retained for independent biochemical validation, rather than as confirmed partners at this stage. Among these candidates, we detected known interactors, including PARP1^32^ and core histones^22^, supporting the specificity of the purification. We also identified the histone variant mH2A1. mH2A, the largest histone, has a tripartite structure comprising an H2A-like histone fold, an unstructured linker, and a non-histone macrodomain^33,34^. The two mH2A isoforms, mH2A1 and mH2A2, are encoded by *H2AFY* and *H2AFY2*, respectively; alternative splicing of *H2AFY* further generates two splice variants, mH2A1.1 and mH2A1.2, which differ in their macrodomain exons^33,34^. Consistent with the recovery of mH2A1, mass spectrometry analysis of histone modifications in purified cGAS-FHA-Nuc also detected the repressive histone mark H3K27me3 with the highest peptide count (Supplementary Table 1), which is known to be enriched on mH2A1-loaded chromatin^35^. Together, these mass spectrometry results identify mH2A1 and H3K27me3 as candidate features of cGAS-associated chromatin, consistent with and extending prior knowledge of mH2A1-chromatin biology.

**Fig. 1.**
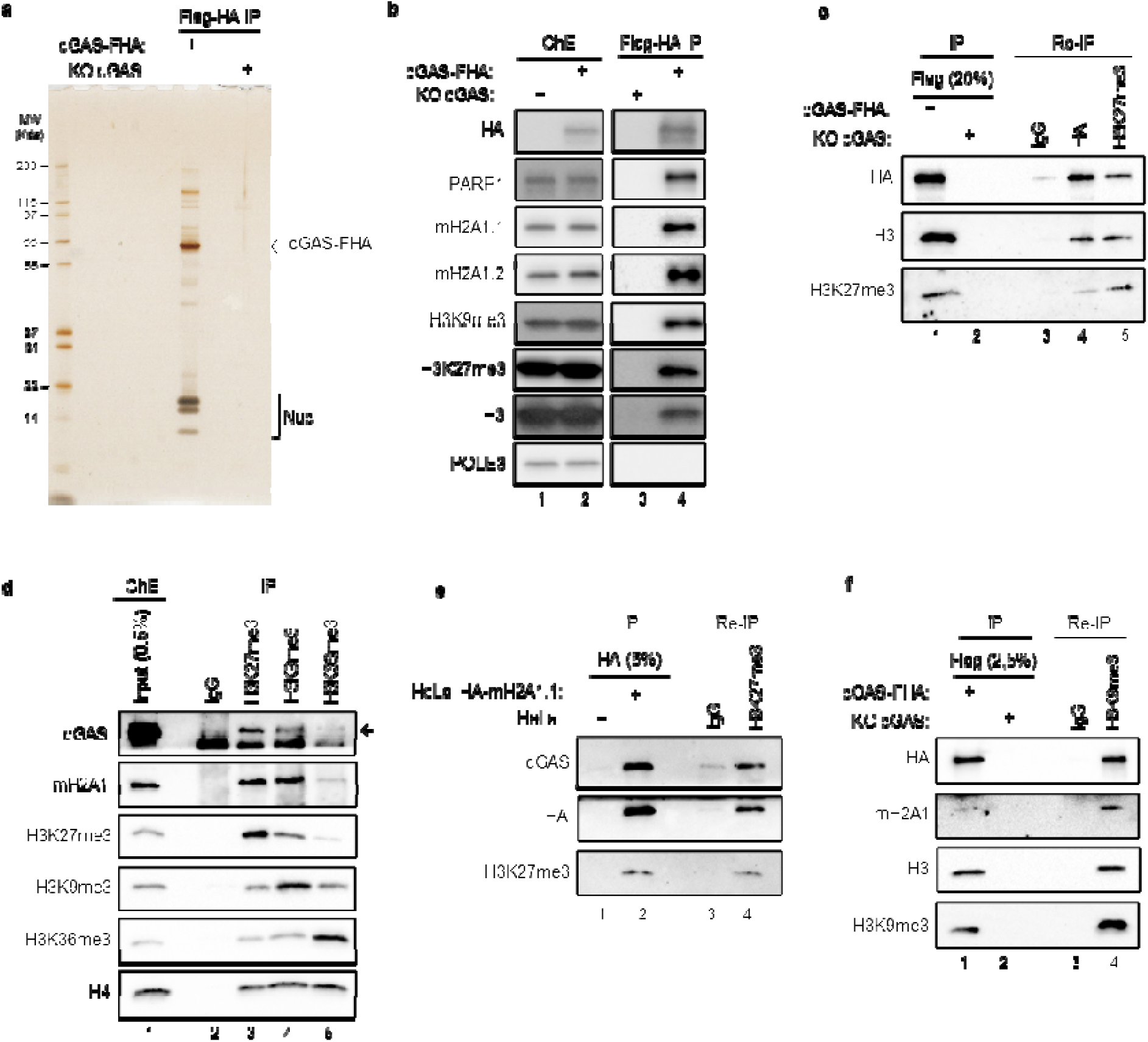
Purification of cGAS from solubilized chromatin reveals binding to mH2A1 and H3K27me3. **a**, FLAG-HA epitope-tagged cGAS (cGAS-FHA) was purified from chromatin extracts (ChE) of cGAS-KO HeLa cells stably expressing cGAS-FHA (lane 1) or empty vector (lane 2). cGAS-FHA was sequentially purified using anti-FLAG and anti-HA antibody-conjugated agarose beads. Proteins were resolved by SDS-PAGE and visualized by silver staining. The identities of cGAS-FHA-associated proteins and histone tail modifications were determined by mass spectrometry (MS) (see also Supplementary Table 1). **b**, The experiment was performed as in (a), except that immunopurified materials were resolved by SDS-PAGE, and the presence of cGAS-FHA-associated proteins and histone tail modifications was assessed by immunoblotting. **c**, Reciprocal immunoprecipitation (RE-IP) performed on cGAS-FHA or KO cGAS ChE. Peptide eluates from FLAG-IPed cGAS-FHA were further subject to Re-IP using HA- or H3K27me3-specific antibody, or rabbit control IgG. Proteins were resolved by SDS-PAGE and analyzed by immunoblotting. **d**, Endogenous cGAS associates with H3K27me3- and H3K9me3-marked chromatin. HeLa ChE was subjected to IP using H3K27me3-, H3K9me3-, H3K36me3-specific antibodies or control rabbit IgG. Immunoprecipitated material was analyzed by immunoblotting using the indicated antibodies. **e**, Re-IP performed on ChE from cells stably expressing HA-mH2A1.1 or untransduced controls. Peptide eluates from HA IP were subject to Re-IP using H3K27me3-specific antibody or rabbit control IgG. **f**, Experiment performed as in (c), using H3K9me3-specific antibodies for Re-IP.

To validate these candidates, we first assessed these interactions by co-immunoprecipitation using ectopically expressed proteins. Western blotting of cGAS-FHA-Nuc immunoprecipitates recovered the known cGAS partner PARP1^32^ and the H3K9me3 mark, previously reported to be enriched on chromatin-bound cGAS^36^. We also confirmed the interaction with mH2A1 and the presence of the H3K27me3 mark, but not of POLE3 protein, used as a negative control (Fig. 1b). The ChE used in these experiments consisted predominantly of mono-nucleosomes (Supplementary Fig. 1c), suggesting that cGAS associates with mono-nucleosomes bearing mH2A1, H3K27me3, and/or H3K9me3. Reciprocally, we derived HeLa cells expressing HA-tagged mH2A1.1 (HA-mH2A1.1) (Supplementary Fig. 1d, compare lane 1 to 2). HA immunoprecipitation (IP) from ChE of HA-mH2A1.1-expressing cells, compared to ChE of untransduced control cells, specifically recovered PARP1, cGAS, H3K27me3, H3K9me3, H4, and HA-mH2A1.1, but not the unrelated HuR protein, which served as a negative control (Supplementary Fig. 1d, compare lane 3 to 4). In order to evaluate the proportion of cGAS bound nucleosomes marked by H3K27me3, we performed reciprocal IPs (Re-IP). Flag-purified cGAS-FHA-Nuc was subjected to a second IP using H3K27me3 antibody or HA antibody which IPs the entire pool of cGAS associated complexes (Fig. 1c). IPs normalized with histone H3 levels show substantial binding of cGAS to H3K27me3 modified nucleosomes (Fig. 1c, compare lanes 4 and 5).

We then sought to confirm these interactions with endogenous proteins. Endogenous cGAS and mH2A1 co-purified with H3K27me3- and H3K9me3-marked nucleosomes but not with IgG or with H3K36me3-marked nucleosomes (Fig. 1d, compare lanes 3 and 4 to lanes 2 and 5), nor with H3K4me3-marked nucleosomes (Supplementary Fig. 1e, compare lane 3 to lanes 2, 4 and 5). Equal H4 levels across IPs indicate that these differences reflect selective enrichment of cGAS and mH2A1 on H3K27me3/H3K9me3-marked nucleosomes, rather than differences in nucleosome recovery efficiency. Collectively, these orthogonal validation experiments, combining ectopic and endogenous co-immunoprecipitation, demonstrate that cGAS-bound nucleosomes are associated with mH2A1, enriched in heterochromatic marks (H3K27me3/H3K9me3), and depleted of H3K36me3 and H3K4me3 (marks of active transcription). Similar results were observed in MRC5 primary human fibroblasts (Supplementary Fig. 1f).

To determine whether cGAS-Nuc partners exist as a single or multiple complexes, we performed glycerol gradient sedimentation of purified cGAS-FHA-Nuc (Supplementary Fig. 1g). Western blotting of gradient fractions revealed cGAS, histones, mH2A1, H3K9me3, H3K27me3, and PARP1 co-sedimenting in fractions 7–9. A small proportion of cGAS was found in lower molecular weight fractions depleted from histones and containing small amounts of PARP1 (Supplementary Fig. 1g, fractions 3–4). This suggests that cGAS-Nuc exists either as a homogeneous complex containing mH2A1 and dual H3K9me3/H3K27me3 marks or as distinct complexes with similar biophysical properties. To resolve this, we performed Re-IPs: HA immunoprecipitation (IP) from ChE of HA-mH2A1.1-expressing cells (Fig. 1e, compare lane 2 to 1) was subjected to Re-IP with H3K27me3 antibody. Substantial amount of cGAS was recovered in H3K27me3-marked nucleosomes containing mH2A1.1 (Fig. 1e, compare lane 4 to 3). Similarly, when Flag-purified cGAS-FHA (Fig. 1f, compare lane 1 to 2) was subjected to ReIP with H3K9me3 antibody, specific enrichment of mH2A1 was observed (Fig. 1f, compare lane 4 to 3) showing that cGAS-FHA-Nuc marked by H3K9me3 also contain mH2A1. Together, these experiments show that cGAS-Nuc complexes can harbor single or dual H3K9me3/H3K27me3 marks alongside mH2A1.

Previous cryo-EM structures reported that cGAS directly binds to the acidic patch of canonical H2A in the nucleosome and is thereby inactivated^23–28^. To reveal the mechanism by which cGAS engages nucleosomes containing mH2A1, we performed cryo-electron microscopy (cryo-EM) analysis. Because mH2A has been reported to preferentially form heterotypic nucleosomes containing one mH2A and one canonical H2A copy in 1:1 stoichiometry^37,38^, we reconstituted heterotypic mH2A1.1-nucleosomes (mH2A1.1Nuc) or mH2A1.2-nucleosomes (mH2A1.2Nuc) by mixing mH2A1.1-H2B or mH2A1.2-H2B dimer, H2A-H2B dimer, H3-H4 tetramer, and 193-bp 601 DNA. Electrophoretic mobility shift assays (EMSA) showed that cGAS bound both mH2A1.1Nuc and mH2A1.2Nuc with higher affinity than the canonical nucleosome (Fig. 2a and 2b), prompting us to determine the structural basis of this preference.

**Fig. 2.**
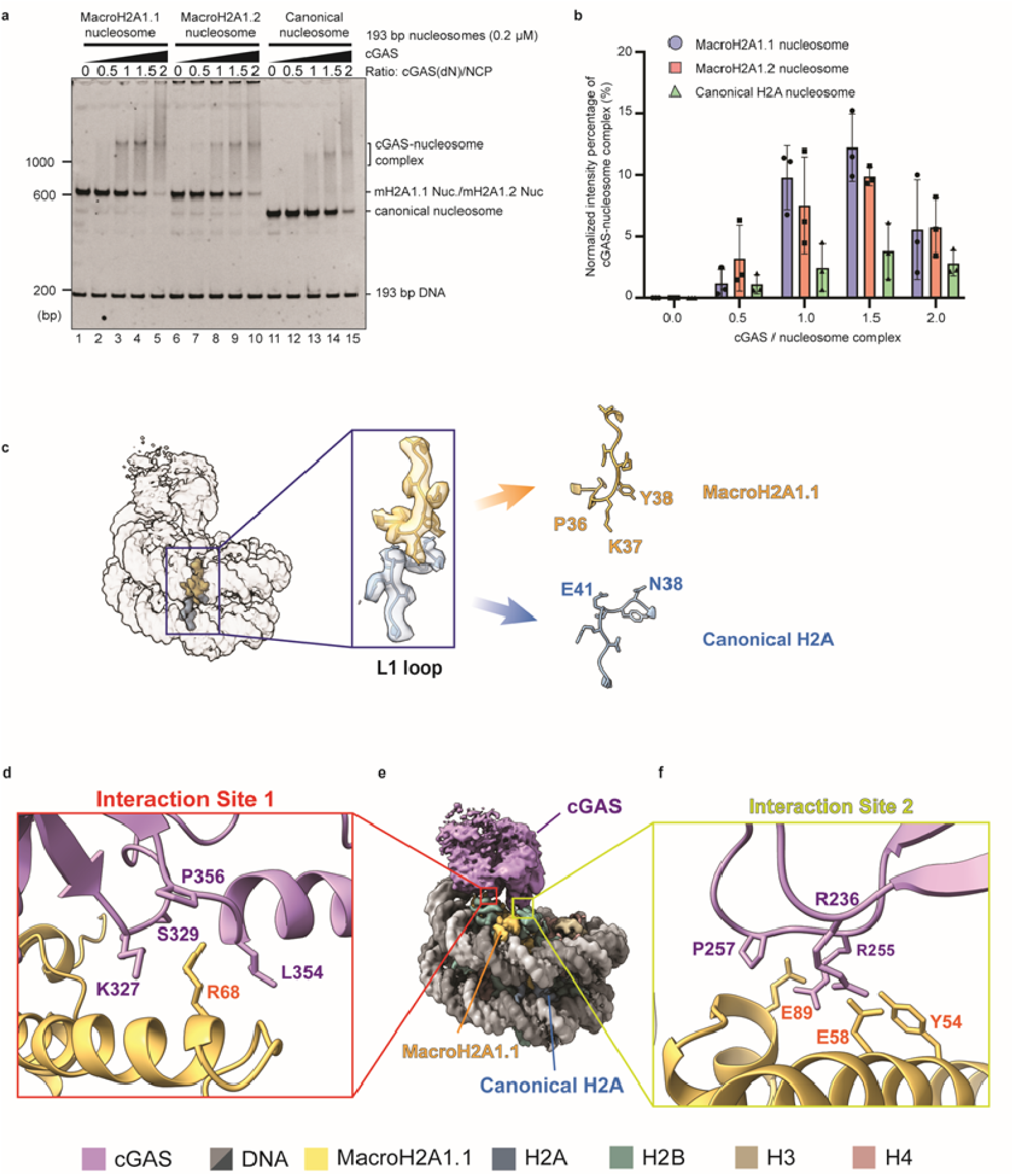
Cryo-EM structure of cGAS bound to mH2A1.1-nucleosome. **a,** Gel electrophoretic mobility shift assay (EMSA) was performed with cGAS and nucleosomes containing mH2A1.1, mH2A1.2, or canonical H2A. Samples were subjected to 4% non-denaturing polyacrylamide gel. The gel was stained with Ethidium Bromide, and bands were detected by a Typhoon imaging analyzer. **b,** Intensities of bands corresponding to unbound nucleosomes were quantitated by Image Quant. Band intensities of each nucleosome without cGAS in each experimental group were referred as 1, estimated cGAS-nucleosome complex was plotted against cGAS/nucleosome ratios (%). Average values of three technical replicates are plotted with ±SD. **c,** Close-up view of the L1 loop region, showing the detailed structures of the L1 loops in mH2A and canonical H2A. **e,** Cryo-EM map of the cGAS-nucleosome complex heterotypically containing mH2A1.1 and canonical H2A. MH2A1.1 (gold), canonical H2A (blue) and cGAS (purple) are displayed. **d, f,** Close-up view of structure model of the cGAS K327, S329, L354, and P356 residues (purple) contacting the mH2A1.1 R68 residue (gold) at the site 1 (d), and the cGAS R236, R255, and P257 residues (purple) contacting the mH2A1.1 Y54, E58, and E89 residues (gold) at the site 2 (f).

To this end, we mixed cGAS with mH2A1.1Nuc, and the resulting cGAS-mH2A1.1Nuc complexes were fractionated by sucrose-gradient ultracentrifugation in the presence of a glutaraldehyde gradient (GraFix)^39^. We determined the structure of the cGAS-mH2A1.1Nuc complex at 2.9 Å resolution (Fig. 2c and Supplementary Fig. 2a-f). This high-resolution structure resolved two distinct L1 loop conformations at the binding regions of canonical H2A and mH2A, allowing us to build an atomic model in which mH2A is unambiguously distinguished from canonical H2A on the basis of L1 loop conformation. The L1 loop interaction within the heterotypic nucleosome is mediated by N38 and E41 of H2A and by P36, K37 and Y38 of macroH2A1.1 (Fig. 2c).

In the cGAS-mH2A1.1Nuc structure, cGAS directly contacts mH2A1.1 within the nucleosome at two distinct interfaces (Fig. 2e). At site 1, cGAS K327, S329, L354 and P356 residues interact with the mH2A1.1 R68 residue (Fig. 2d). At site 2, cGAS R236, R255 and P257 residues contact the mH2A1.1 Y54, E58 and E89 residues, which together form the acidic patch on the nucleosome surface (Fig. 2f). These cGAS-mH2A1.1 contacts closely resemble the previously described cGAS interactions with canonical H2A in the nucleosome^23–28^. Strikingly, however, the cryo-EM density corresponding to cGAS was predominantly detected on the acidic patch of the mH2A1.1 copy, and not on that of the canonical H2A copy, within the same heterotypic nucleosome (Fig. 2e). This asymmetric occupancy indicates that, when given the choice within a single nucleosome, cGAS preferentially engages the mH2A1.1-containing acidic patch over the canonical H2A-containing one.

Next, we assessed the contribution of mH2A1 and H3K27me3 to cGAS binding to chromatin and their impact on cGAS-mediated induction of the type I IFN response. HeLa cells were transfected with non-targeting (NT) control, mH2A1.1-, or mH2A1.2-specific siRNA, and 48 hrs later total, cytoplasmic, and chromatin fractions were prepared. Total extracts were used to analyze the knockdown (KD) efficiency and specificity of the siRNAs used, and their impact on cGAS expression (Fig. 3a). cGAS was extracted from the chromatin fraction using sequentially increasing concentrations of NaCl (Fig. 3b). Consistent with a previous report²¹, western blot analysis using a cGAS-specific antibody showed that cGAS is tightly tethered to chromatin, with maximum elution observed at 0.75 M NaCl in both control and mH2A1.2-KD cells (Fig. 3b, upper panels). Remarkably, KD of mH2A1.1 resulted in maximum solubilization of cGAS at 0.25 M NaCl (Fig. 3b, upper panels). As a control, KD of mH2A1.1 and mH2A1.2 had no effect on H3 binding to chromatin (Fig. 3b, lower panels). Quantification of cGAS solubilization from four independent experiments confirmed the effect of mH2A1.1 KD on cGAS binding to chromatin (Fig. 3c). To address the involvement of H3K27me3 in cGAS tethering to chromatin, similar experiments were performed using UNC1999, an inhibitor of EZH1 and EZH2, which are responsible for H3K27me3 deposition. UNC1999 treatment of HeLa cells reduced total H3K27me3 levels, without affecting cGAS levels (Fig. 3d) and significantly enhanced the amount of cGAS extracted at 0.25 M NaCl, without affecting H3 extraction (Fig. 3e, upper and lower panels, respectively). Quantification of cGAS solubilization from three independent experiments confirmed the effect of UNC1999 on cGAS binding to chromatin (Fig. 3f). Taken together, these experiments highlight the role of mH2A1.1 and H3K27me3 in regulating cGAS tethering to chromatin.

**Fig. 3.**
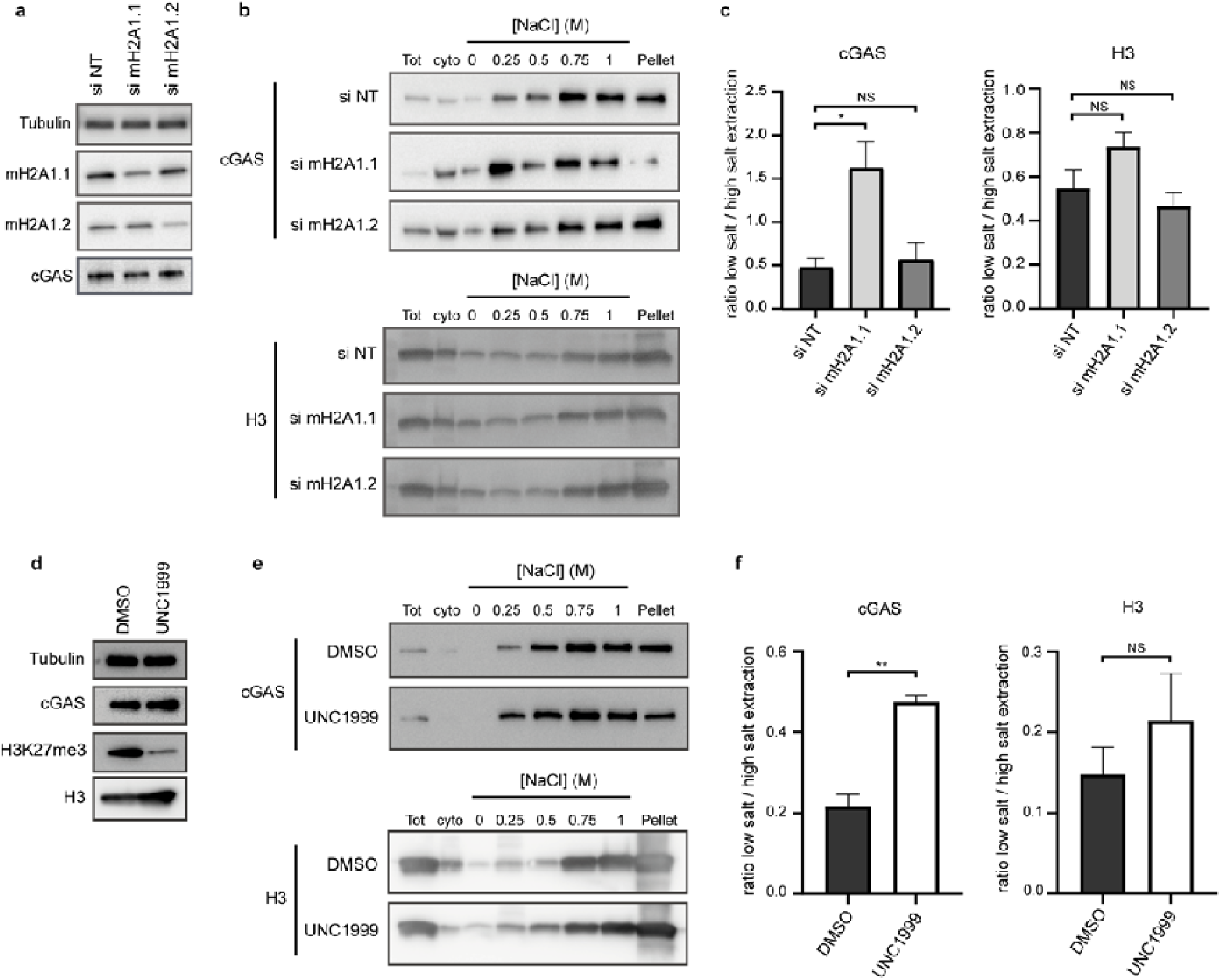
H3K27me3 and mH2A1.1 promote cGAS tethering to chromatin. **a,** HeLa cells were transfected with mH2A1.1-, mH2A1.2-, or non-targeting (NT) control siRNA for 48 h. Depletion of mH2A1.1 and mH2A1.2 was assessed by western blot. **b,** Cells were separated into cytosolic and nuclear fractions, followed by sequential stepwise elutions of nuclear pellets with the indicated concentrations of NaCl. cGAS and H3 extraction was monitored by western blot. One representative experiment of four independent experiments is shown. **c,** Ratios of cGAS and H3 in low-salt (0.25 M) vs. high-salt (1 M) extraction were determined by densitometry. Mean ± SEM of three independent experiments. **d,** HeLa cells were treated with DMSO or 5 µM UNC1999 for 48 h. Total levels of H3K27me3 were assessed by western blot. **e,** Cells were separated as in (b). cGAS and H3 extraction was monitored by western blot. One representative experiment of three independent experiments is shown. **f,** Ratios of cGAS and H3 in low-salt (0.25 M) vs. high-salt (1 M) extraction were determined by densitometry. Mean ± SEM of three independent experiments. *P < 0.05, **P < 0.01, ***P < 0.001; independent Student’s t-test.

We then analyzed the role of mH2A1 and H3K27me3 in regulating cGAS-mediated activation of the cGAS-STING-IRF3 innate immune pathway. HeLa cells were treated with DMSO, the cGAS inhibitor G140, or the STING inhibitor H151 for 3 hrs before transfection with NT, mH2A1.1-, or mH2A1.2-specific siRNAs. Cells were subsequently harvested for protein and RNA analysis. RNA was used to quantify interferon-β (IFNβ) and interferon-stimulated genes (ISGs) by quantitative RT-PCR. As shown in Fig. 4a, KD of mH2A1.1, but not of mH2A1.2, resulted in a cGAS- and STING-dependent induction of ISG15, IFIT1, and MXA mRNAs, as well as IFNβ, IFIT2, MX2 and RSAD2 mRNAs (Supplementary Fig. 3a). This induction of ISGs upon mH2A1.1 KD was confirmed using additional siRNAs (Supplementary Fig. 3b). Western blot analyses showed that STING and IRF3 phosphorylation were increased upon KD of mH2A1.1 but not of mH2A1.2 (Fig. 4b, left panel). Both G140 and H151 prevented or decreased STING and IRF3 phosphorylation in mH2A1.1-KD HeLa cells (Fig. 4b, right panel).

**Fig. 4.**
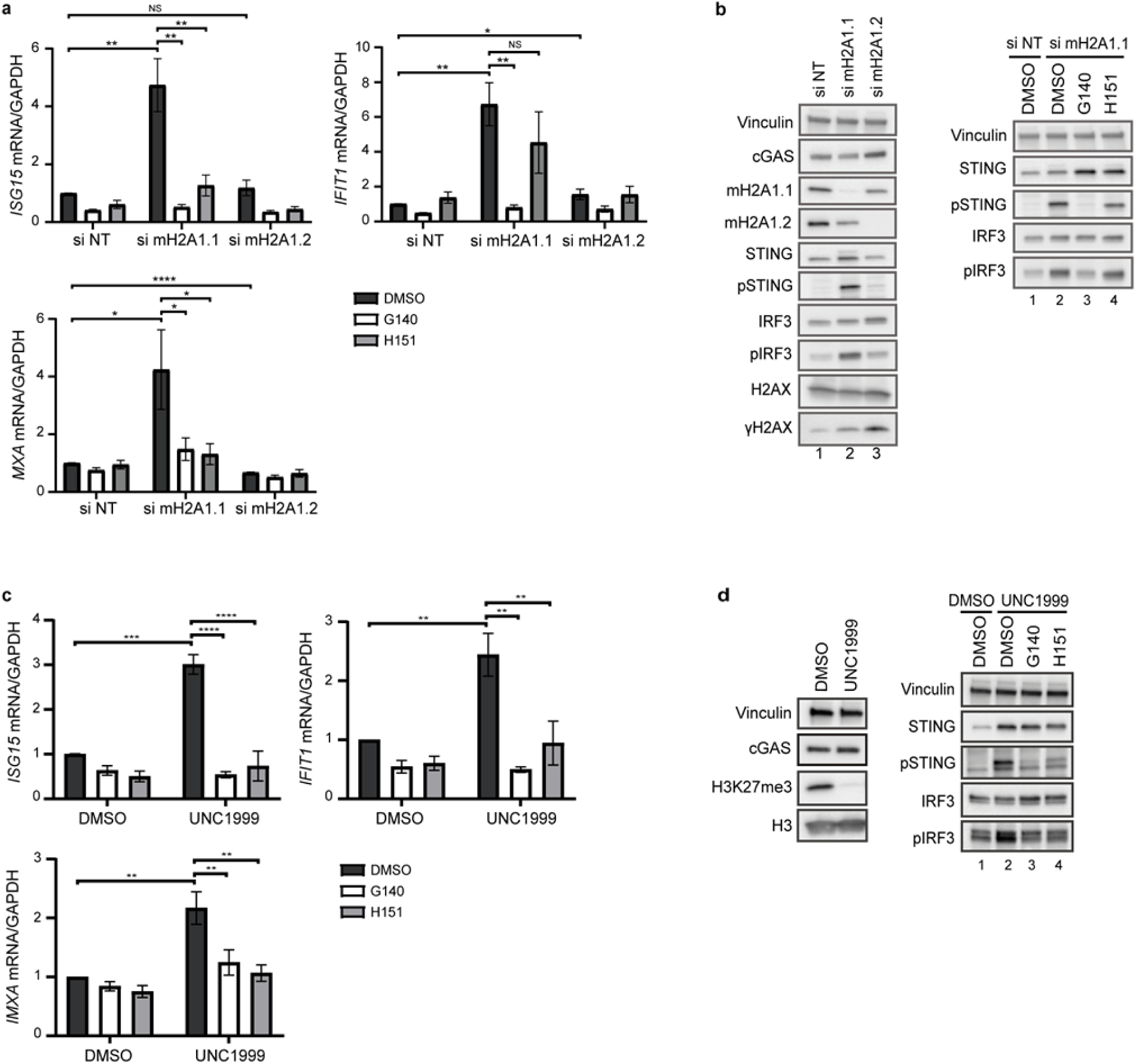
Depletion of H3K27me3 and mH2A1.1 results in cGAS-dependent innate immune activation. **a,** HeLa cells were transfected with mH2A1.1-, mH2A1.2-, or non-targeting (NT) control siRNA for 72 h. ISG15, IFIT1, and MXA mRNA expression were measured by RT-qPCR (normalized to GAPDH mRNA). Mean ± SD of four independent experiments. **b,** Western blot analysis of the canonical cGAS activation pathway upon mH2A1.1 and mH2A1.2 siRNA transfection. One representative experiment of four independent experiments is shown. **c,** HeLa cells were treated with DMSO or 5 µM UNC1999 for 48 h. ISG15, IFIT1, and MXA mRNA expression were measured by RT-qPCR (normalized to GAPDH mRNA). Mean ± SD of four independent experiments. **d,** Western blot analysis of the canonical cGAS activation pathway upon UNC1999 treatment. One representative experiment of four independent experiments is shown. *P < 0.05, **P < 0.01, ***P < 0.001; independent Student’s t-test.

Notably, the induction of the cGAS-STING pathway upon mH2A1.1 KD is independent of the DNA damage response, as measured by γH2AX levels. Indeed, while higher levels of γH2AX were observed upon KD of mH2A1.2 compared to mH2A1.1, no activation of the IFN pathway was observed in mH2A1.2-KD cells (Fig. 4b, left panel, compare lane 3 to lane 2). Furthermore, although both mH2A1.1 siRNAs induced the cGAS-STING pathway, only siRNA mH2A1.1#2 induced γH2AX (Supplementary Fig 3b), further dissociating cGAS-STING activation from DNA damage signaling.

Similarly, treatment with UNC1999 resulted in a cGAS-/STING-dependent induction of ISG15, IFIT1 and MXA mRNAs (Fig. 4c), as well as IFNβ, IFIT2, MX2 and RSAD2 mRNAs (Supplementary Fig. 3c). Western blot analyses showed that UNC1999 efficiently reduced H3K27me3 levels in treated HeLa cells (Fig. 4d, left panel) and enhanced STING and IRF3 phosphorylation in a cGAS-and STING-dependent manner (Fig. 4d, right panel). Taken together, our data identify mH2A1.1 and H3K27me3 as regulators of cGAS tethering to chromatin, whose depletion leads to induction of an innate immune response in HeLa cells, independently of DNA damage signaling.

The biochemical interactions of cGAS with mH2A1 and heterochromatin marks prompted us to inspect its genomic distribution alongside that of mH2A1, H3K27me3, and H3K9me3. To this end, the genomic distribution of cGAS-FHA, mH2A1.1, mH2A1.2, H3K27me3, and H3K9me3 was analyzed using EV or cGAS-FHA reconstituted cGAS KO HeLa cells. Additionally, cGAS, mH2A1.1, and mH2A1.2 genomic distribution were also analyzed in cells transfected with their respective specific siRNAs (Fig. 5a. Supplementary Fig. 4 and supplementary Fig. 5a). Chromatin immunoprecipitation followed by high-throughput sequencing (ChIP-seq) was performed using specific antibodies, after validating their specificity by quantitative PCR using specific primers corresponding to positive and negative control genomic regions (Supplementary Fig. 4). cGAS ChIP-seq revealed the presence of significant (> 5,000) specific cGAS peaks (Fig. 5a) validated by their loss in cGAS KO cells (Supplementary Fig. 5a). Analysis of epigenetic mark levels associated with such cGAS peaks showed low levels of gene-associated euchromatic marks (H3K4me3 and H3K36me3) compared to random control peaks, contrasting with significantly higher levels of heterochromatic marks (H3K9me3 and H3K27me3) and DNA methylation (Fig. 5a, b). Reduced levels of DNase sensitivity, active-gene-associated H3K36me3, and enhancer-associated H3K27 acetylation further suggested that cGAS binds to regions outside active genes and enhancers, typically associated with active chromatin regions (Fig. 5b). To further inspect cGAS peaks beyond the total averaged peaks, a clustering approach was performed using the factors found to interact with cGAS, namely mH2A1.1, mH2A1.2, H3K9me3, and H3K27me3 marks (Fig. 5c and Supplementary Fig. 5b). Eight distinct clusters were identified, distinguishable by their H3K9me3-to-H3K27me3 ratio, the “oriented” profile of cGAS peaks (clusters 6–8), the absence of such orientation (central peaks in clusters 2, 3, and 5), or signal dispersion into domain-like distributions involving neighboring peaks (clusters 1 and 4). All cGAS peaks were enriched in mH2A1 variants (Fig. 5c and Supplementary Fig. 5b). All clusters except cluster 4 were enriched in TSSs and/or CTCF sites (Fig. 5d, e). Notably, cluster 4 was particularly enriched in centromeric repeat sequences (77% of such sites) (Fig 5e), confirmed by its high levels in CENP-A and γ-H2AX (supplementary Fig. 5c). Cluster 4 (14% of total cGAS peaks) aligns with prior reports of cGAS-centromere association^36^, but we also identified thousands of additional heterochromatic cGAS-binding sites with elevated H3K27me3 relative to H3K9me3 (clusters 5 to 8). Analyses of cGAS peaks enrichment in Encode TF IDR peaks (Encode) revealed relative enrichment in CTCF, Rad21 and SMC3 (cohesion complex subunits) at clusters 6-8 (Fig. 5e and Supplementary Fig. 5d). CTCF is an architectural protein, which plays an important role in 3D chromatin organization and transcriptional regulation^40^. Over the genome, CTCF binding sites can be defined as low or high occupancy depending on their affinity for CTCF. High occupancy sites are involved in 3D genome organization and are mostly associated with high insulation of neighboring regions while low occupancy sites are involved in transcriptional regulation, and display little to no effect on the overall insulation profile of these regions^40^. The CTCF binding levels across cGAS bound regions were generally lower than those of randomly selected CTCF peaks (Fig. 5f), suggesting that CTCF sites across cGAS bound regions could correspond to low-occupancy sites with low insulation function (Supplementary Fig. 5e-f). In fact, selecting CTCF peaks to match (CTCF_MATCH) the levels observed within cGAS bound regions appears to select for mostly CTCF peaks with little to no effect on the overall insulation profile of these regions (Supplementary Fig. 5e-f). Unexpectedly, inspecting the overall genomic insulation levels across cGAS bound regions revealed that cGAS regions overlapping CTCF peaks showed a clear linkage to TAD boundaries (Fig. 5g and Supplementary Fig. 5f), with significantly higher insulation levels than CTCF_MATCH and random cGAS peaks (cGAS_RANDOM), and comparable insulation levels of random CTCF peaks (CTCF_RANDOM) (Fig. 5g and Supplementary Fig. 5e-f). This implies that cGAS-CTCF-overlapping peaks possess unique chromatin features related to TAD borders. Supporting this, mH2A1 was exclusively associated with CTCF peaks bound by cGAS (Supplementary Fig. 6a,b), which were highly marked by H3K27me3 compared to cGAS-negative, mH2A1-unbound CTCF peaks (Supplementary Fig. 6c).

**Fig. 5.**
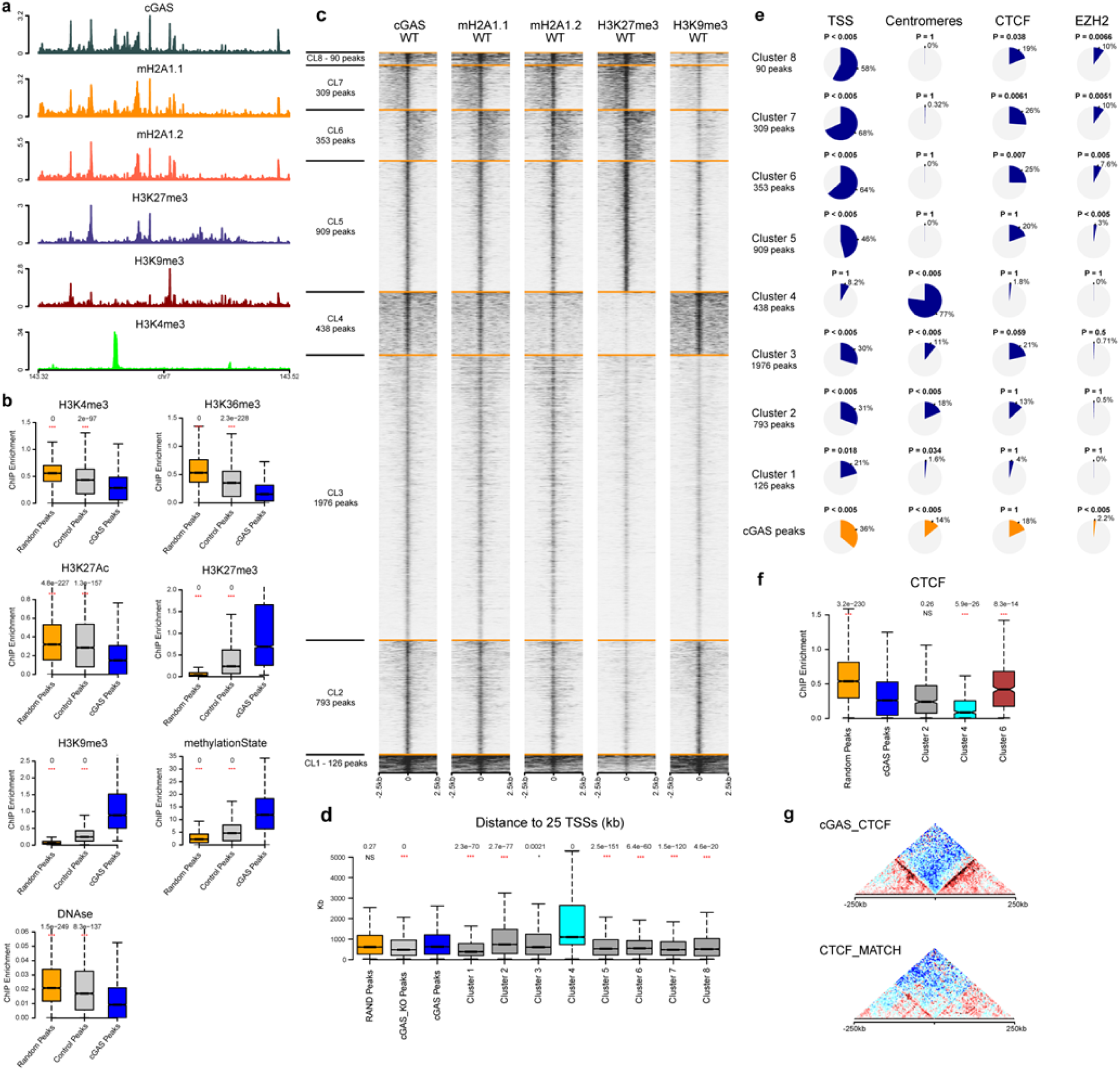
cGAS co-localizes with mH2A1.1/2 histone variants and H3K27me3 or H3K9me3 epigenetic marks. **a**, Visualization of a 200-kb region on chromosome 17 illustrating ChIP-seq data obtained in HeLa cells for cGAS, mH2A1.1, mH2A1.2, H3K27me3, H3K9me3 (this study), and H3K4me3 (ENCODE). **b**, Comparison of ChIP-seq enrichment levels across cGAS binding sites, random genomic loci, and control loci (peaks identified in cGAS-KO samples) for histone marks associated with euchromatin (H3K4me3, H3K36me3), H3K27 state modifications (acetylation or methylation), constitutive heterochromatin marker (H3K9me3), DNA methylation percentage, and chromatin accessibility (DNase). P values were computed using a two-sided Wilcoxon test against cGAS peaks (blue). **c**, K-means clustering of cGAS-identified peaks based on 2.5-kb regions surrounding each peak (50-bp resolution) for cGAS, mH2A1.1, mH2A1.2, H3K27me3, and H3K9me3 profiles. cGAS predominantly co-localizes with mH2A1.1/2 and exhibits variable levels of H3K27me3 and H3K9me3, as well as binding patterns (centered, asymmetric, dispersed). **d**, Box plots showing linear genomic distance distributions of each cGAS epigenetic cluster to the nearest 25 annotated transcription start sites (TSSs). Cluster 4 exhibits significantly greater distances, consistent with peaks mapping to gene deserts. P values were derived by comparing each cluster’s distribution to all cGAS peaks (blue) using a two-sided Wilcoxon test**. e**, Pie charts displaying the proportion of cGAS peaks in each cluster overlapping annotated TSSs, centromeres, CTCF peaks, or EZH2 peaks. P values were calculated via a permutation test (200 permutations) compared to randomized genomic coordinates, with an overlap threshold of 1 kb. **f**, CTCF ChIP-seq enrichment scores across random peaks, all cGAS peaks, and cGAS peaks in epigenetic clusters 2, 4, and 6. P values were determined using a two-sided Wilcoxon test comparing each cluster to all cGAS peaks (blue). **g**, Aggregate Micro-C maps across cGAS peaks overlapping CTCF sites (upper two-dimensional [2D] plot) and CTCF peaks with no cGAS but matched enrichment levels (CTCF_MATCH, lower 2D plot) (see Methods and Supplementary Fig. 5e for quantification).

Next, we inspected the interplay between cGAS and epigenetic marks H3K9me3 and H3K27me3, and the histone variants mH2A1.1 or mH2A1.2. To this end, we re-quantified the levels of these factors after depleting cGAS, mH2A1.1, or mH2A1.2 (Fig. 6a; Supplementary Fig. 7a-c). Strikingly, cGAS depletion reduced mH2A1.1 levels at all clusters except the centromeric cGAS cluster (cluster 4), while mH2A1.1 depletion similarly affected cGAS levels (Fig. 6a,b; Supplementary Fig. 7a-c). Additionally, H3K9me3 levels increased at all cGAS sites upon depletion of cGAS, mH2A1.1, or mH2A1.2 (Fig. 6a,c; Supplementary Fig. 7a-c). cGAS depletion also decreased H3K27me3 levels at CTCF-, cohesin-, and TSS-enriched sites (Supplementary Fig. 7d), whereas mH2A1.1 or mH2A1.2 depletion elevated H3K27me3 levels (Fig. 6a and Supplementary Fig. 7a–b). These results confirm the biochemical interplay between mH2A1.1 and cGAS and/or H3K27me3, alongside reduced centromere-associated constitutive heterochromatin mark H3K9me3 (cluster 4). Overall, chromatin dynamics following cGAS, mH2A1.1, or mH2A1.2 depletion can be categorized into three major patterns (Fig. 6c). First, centromeric peaks (cluster 4) showed no mutual influence between mH2A1.1 and cGAS (Fig. 6d, left panel and Supplementary Fig. 7a, b). Second, epigenetic clusters 5–8 exhibited strong interdependence between cGAS and mH2A1.1, with increased H3K27me3 levels upon mH2A1.1 or mH2A1.2 depletion (Fig. 6d, middle panel and Supplementary Fig. 7a, b). Third, epigenetic clusters 2 and 3 displayed cGAS-mH2A1.1 interdependence and elevated H3K9me3 levels after cGAS or mH2A1.1 depletion (Fig. 6d, right panel; Supplementary Fig. 7a,b). These cGAS peak subsets retained centromere, TSS, CTCF, and EZH2 binding enrichment profiles matching their epigenetic clusters (Supplementary Fig. 7d). Importantly, no cGAS clusters showed interdependence between cGAS and mH2A1.2, consistent with mH2A1.2 KD having no effect on cGAS-chromatin tethering (Fig. 3b–c).

**Fig. 6.**
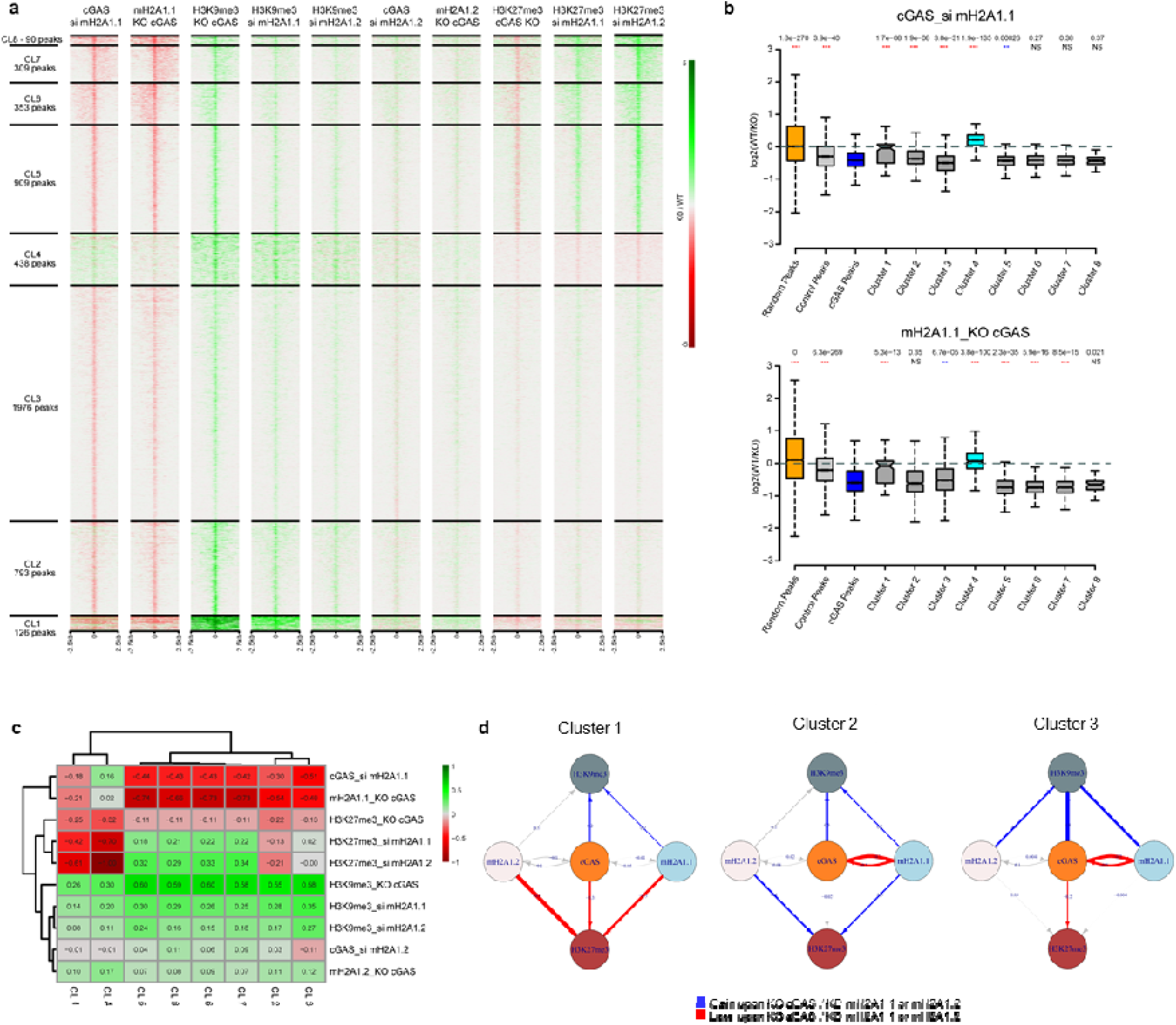
Synergistic binding of cGAS and mH2A1.1 differentially regulates H3K27me3 and H3K9me3 levels depending on peak localization. **a**, Differential ChIP-seq signal (KO vs. WT) assessing the effect of cGAS, mH2A1.1-, or mH2A1.2 depletion on cGAS, mH2A1.1, mH2A1.2, H3K27me3, and H3K9me3 levels across predefined cGAS peak clusters (Fig. 5c). **b**, Quantification of differential ChIP-seq signal (log2(WT/KO)) distribution for each cGAS peak cluster. Top: cGAS level variation upon mH2A1.1 siRNA depletion. Bottom: mH2A1.1 level variation in cGAS-KO HeLa cells. P values were determined by comparing each cluster’s distribution to all cGAS peaks (blue) using a two-sided Wilcoxon test. **c**, Hierarchical clustering analysis of average ChIP-seq signal variation (log2(KO/WT)) across cGAS epigenetic clusters for cGAS, mH2A1.1, mH2A1.2, H3K27me3, and H3K9me3, highlighting three major behaviors: Centromere-specific (clusters 1 and 4). H3K27me3 gain upon cGAS or mH2A1.1 depletion (clusters 5–8). Low H3K27me3 levels (clusters 2 and 3). **d**, Graphical summary of the mutual regulatory influence between cGAS and mH2A1.1 (compared to mH2A1.2) on H3K27me3 and H3K9me3 levels, categorized by cluster type (centromeric, H3K27me3 gain, or low H3K27me3).

## Discussion

The DNA sensor cGAS is paradoxically localized in the nucleus, bound to chromatin in an enzymatically inactive state^20,22,29^. Prior structural studies established that cGAS binds nucleosomes via the H2A/H2B acidic patch to prevent aberrant immune activation by self-DNA^23–28^; however, these studies relied on nucleosomes reconstituted in vitro from canonical histones alone, a system that could not, by design, reveal a preference for non-canonical histone variants. This mode of inhibition has consequently been described as compartment-agnostic: any canonical nucleosome was thought capable of sequestering cGAS, largely irrespective of the chromatin state in which it resides. By contrast, our biochemical purification of native cGAS-bound nucleosomes from cells shows that the nucleosomes cGAS actually engages in vivo are not canonical, but heterotypic particles containing the histone variant mH2A1. A recent comprehensive review of the cGAS-STING pathway further noted that nuclear cGAS is constrained by additional layers of control, including competition with barrier-to-autointegration factor (BAF) for DNA access, restriction of linker DNA exposure by histone H1, and proteostatic turnover via the SPSB3-CRL5 ubiquitin ligase complex, and explicitly identified how nucleosomal regulation of cGAS integrates with these mechanisms as an important open question^41^. Our work addresses this question directly: rather than being sequestered indiscriminately by any nucleosome, cGAS is selectively directed toward mH2A1- and H3K27me3-marked chromatin, adding histone variant- and modification-dependent targeting as a spatially resolved layer within this regulatory network. Three lines of evidence support this conclusion. First, cGAS-bound nucleosomes purified from cells are selectively enriched for mH2A1 and for the heterochromatic marks H3K27me3 and H3K9me3. Second, cryo-EM structures of cGAS bound to heterotypic nucleosomes reveal that cGAS preferentially occupies the acidic patch on the mH2A1 side rather than the canonical H2A side of the same particle, providing a structural basis for selective recruitment to mH2A1-containing chromatin rather than a generic preference for any nucleosome. Third, depleting mH2A1.1 or pharmacologically blocking H3K27me3 deposition releases cGAS from chromatin and triggers cGAS- and STING-dependent induction of interferon-stimulated genes, independently of concurrent DNA damage signaling. Together, these findings reframe nucleosomal sequestration of cGAS not as a uniform, chromatin-state-independent lock, but as a targeted mechanism that couples cGAS storage to facultative heterochromatin.

Our structural data raise a specific mechanistic question that remains open. The cGAS residues R236, R255, and P257 contact the mH2A1.1 acidic patch residues Y54, E58, and E89, positions that are conserved in canonical H2A, yet cGAS density is asymmetrically enriched on the mH2A1.1 side of the heterotypic nucleosome. Because the mH2A macrodomain is flexible and not resolved in our cryo-EM map, we cannot yet distinguish between two non-exclusive possibilities: the macrodomain may sterically favor cGAS engagement on the mH2A1.1 side, for instance by restricting access or altering local nucleosome dynamics on the canonical H2A side, or unresolved differences elsewhere in the mH2A1.1 histone fold may enhance acidic patch accessibility or affinity independently of the macrodomain. Distinguishing these scenarios will require structural and biochemical dissection of macrodomain-truncated or acidic patch-swapped nucleosomes, a direction we identify as a priority for future work.

Our genome-wide analyses further reveals that cGAS occupies two epigenetically and functionally distinct chromatin compartments. The majority of cGAS peaks reside at chromosome-arm sites co-occupied by mH2A1.1, H3K27me3, and CTCF, where cGAS and mH2A1.1 binding are mutually interdependent. Loss of either cGAS or mH2A1.1 increases H3K9me3 at these sites, suggesting a dynamic competition between facultative and constitutive heterochromatin at loci normally stabilized by cGAS-mH2A1.1 co-binding. By contrast, centromeric cGAS clusters (14% of peaks) are independent of mH2A1.1, enriched in H3K9me3, γH2AX, and CENP-A, and consistent with prior reports of cGAS-centromere association^36^. This mH2A1.1-independence at centromeres may reflect a context-specific anchoring mechanism that substitutes for mH2A1.1 in these regions. That two mechanistically distinct modes of chromatin tethering converge on the same protein, cGAS, underscores that nuclear chromatin environments deploy specialized, locally adapted strategies to control its activity rather than a single universal mechanism.

The association we observe between cGAS-bound CTCF sites and TAD boundary insulation adds a further, unanticipated dimension to this picture. cGAS-occupied CTCF sites display insulation levels comparable to canonical high-occupancy CTCF peaks despite showing lower CTCF occupancy themselves, suggesting that cGAS binding, or the chromatin state that recruits cGAS, may be associated with boundary elements that acquire insulating function through a mechanism distinct from CTCF occupancy alone. This observation is correlative, and we do not yet know whether cGAS is a passive marker of these boundaries, a structural contributor to their insulation, or a bystander recruited by the same chromatin features that establish them. Supporting the plausibility of a direct functional link, a recent study in regulatory T cells demonstrated that nuclear cGAS physically interacts with CTCF through its central zinc-finger DNA-binding domain, and that this cGAS-CTCF interaction is required to sustain CTCF binding and chromatin looping at the *Tnfrsf9* and *Tnfrsf4* loci, independently of cGAS catalytic activity or STING signaling^42^. This precedent establishes that cGAS can directly modulate CTCF-dependent chromatin architecture in at least one cellular context. The cGAS-CTCF association we observe genome-wide in HeLa cells may thus reflect a similar direct interaction rather than mere co-occurrence. This finding would represent a starting point for future investigation of 3D genome organization rather than as an established causal role for cGAS in TAD boundary formation.

Beyond its mechanistic interest, our findings carry implications for the physiological and pathological consequences of nuclear cGAS regulation. Supporting the pathological relevance of dysregulated chromatin-associated cGAS activation, a recent study demonstrated that unrepaired mitotic DNA-protein crosslinks trigger micronuclei formation and cGAS-STING-dependent innate immune activation in mice, driving premature aging phenotypes and embryonic lethality that were substantially rescued by genetic or pharmacological cGAS-STING inhibition^43^. Together with our findings, this suggests that chromatin-derived cGAS activation, whether triggered by loss of nucleosomal tethering, as we describe here, or by structural DNA lesions that generate cGAS-exposing micronuclei, converges on a shared, therapeutically tractable innate immune axis. The convergence of cGAS and mH2A1 on genome stability, senescence, DNA damage response, and telomere maintenance^15,32,44–53^ further suggests that the mH2A1-H3K27me3-cGAS tethering axis we describe may be relevant not only to acute innate immune activation but also to the chronic, low-grade inflammatory states associated with cellular senescence and aging.

This axis may also help explain a long-standing observation in tumor immunology. Pharmacological EZH1/EZH2 inhibitors, already in clinical development to reprogram immunologically “cold” tumors (immune-desert) into “hot,” T cell-infiltrated ones^54–56^, have been reported to reactivate type I interferon signaling in cancer cells^57^, an effect generally attributed to derepression of chemokine (for example, CXCL9/CXCL10) and antigen-presentation genes^58–60^. Our data raise the possibility that this response is driven, at least in part, by direct release of cGAS from H3K27me3-marked chromatin rather than, or in addition to, transcriptional derepression of individual target genes, positioning the mH2A1-H3K27me3-cGAS tethering axis as a chromatin-intrinsic determinant of tumor immunogenicity and suggesting that EZH1/EZH2 inhibitors may act in part as indirect cGAS-STING agonists. Modulating this axis more broadly, for example by targeting mH2A1.1 loading, H3K27me3 deposition, or the cGAS-mH2A1.1 interface directly, may therefore offer a chromatin-directed strategy for tuning cGAS-STING signaling across disease contexts ranging from autoinflammatory disorders and aging to cancer, complementing existing efforts targeting cGAS catalytic activity or STING trafficking directly.

## Acknowledgements

We wish to thank members of the Molecular Virology lab, S. Khochbin and D. Trono for critical reading of the manuscript and for helpful discussion. We are grateful to S.Abbou for providing valuable technical assistance. We also thank Y. Iikura, Y. Takeda, and M. Dacher (The University of Tokyo) for their assistance. A. G. Ladurner and M. Buschbeck kindly provided the mH2A1.1 and mH2A1.2 expression plasmids. This work was supported by grants from European Research Council ERC-2018-ADG (RetroChrom#835184), and ANRS-MIE to M.B. (S.T-H) was supported by ERC-2018-ADG (RetroChrom#835184) and ANRS fellowship. C.J. was supported by MERT and ANRS-MIE fellowship. The work was supported in part by ANRS (LS229035) and by MSD-Avenir to O.C. This work was supported in part by JSPS KAKENHI Grant Numbers JP22K18034 [to S.M.], JP22K15033 [to T.K.], JP23K17392 [to T.K.], JP24H00062 [to T.K.], JP23H05475 [to H.K.], JP24H02328 [to H.K.], and JP24H02319 [to H.K.]; JST PRESTO Grant Number JPMJPR2288 [to S.M.], JST ERATO Grant Number JPMJER1901 [to H.K.], JST CREST Grant Number JPMJCR24T3 [to H.K.], and Research Support Project for Life Science and Drug Discovery (BINDS) from AMED under Grant Numbers JP25ama121009 [to H.K.].

## Author contribution

M.B.; S.B. and H.K. conceived the study. All authors co-designed experiments and interpreted data. T.H.S.; C.J.; B.S.; W.S.; L.B.M. and S.M. performed experiments. G.P. and O.C. performed the analysis and statistics of the genomic data. M.B. and H.K. wrote the manuscript. All authors commented and edited the manuscript.

## Data availability

All data are available in the main text or the supplementary materials. All reagents are available upon reasonable request to the corresponding author.

## Supplementary Materials

Materials and Methods

Figs. S1 to S7

Tables S1

References (61–77)

## References

1. Stratigi, K., Chatzidoukaki, O. & Garinis, G. A. DNA damage-induced inflammation and nuclear architecture. Mech. Ageing Dev. 165, 17–26 (2017).

2. Nihira, N. T., Kudo, R. & Ohta, T. Inflammation and tumor immune escape in response to DNA damage. Semin. Cancer Biol. 110, 36–45 (2025).

3. Goddard, A. M., Cho, M.-G., Lerner, L. M. & Gupta, G. P. Mechanisms of Immune Sensing of DNA Damage. J. Mol. Biol. 436, 168424 (2024).

4. Hayman, T. J. & Glazer, P. M. Regulation of the Cell-Intrinsic DNA Damage Response by the Innate Immune Machinery. Int. J. Mol. Sci. 22, 12761 (2021).

5. Ragu, S., Matos-Rodrigues, G. & Lopez, B. S. Replication Stress, DNA Damage, Inflammatory Cytokines and Innate Immune Response. Genes 11, 409 (2020).

6. Duzanic, F. D. & Penengo, L. The interferon response at the intersection of genome integrity and innate immunity. DNA Repair 145, 103786 (2025).

7. Tong, J. et al. When DNA-damage responses meet innate and adaptive immunity. Cell. Mol. Life Sci. CMLS 81, 185 (2024).

8. Du, Q., Miao, Y., He, W. & Zheng, H. ADP-Ribosylation in Antiviral Innate Immune Response. Pathogens 12, 303 (2023).

9. Brégnard, C., Benkirane, M. & Laguette, N. DNA damage repair machinery and HIV escape from innate immune sensing. Front. Microbiol. 5, 176 (2014).

10. Ma, Y. et al. Growth arrest and DNA damage-inducible 45: a new player on inflammatory diseases. Front. Immunol. 16, 1513069 (2025).

11. Morchikh, M. et al. HEXIM1 and NEAT1 Long Non-coding RNA Form a Multi-subunit Complex that Regulates DNA-Mediated Innate Immune Response. Mol. Cell 67, 387–399.e5 (2017).

12. Laguette, N. et al. Premature activation of the SLX4 complex by Vpr promotes G2/M arrest and escape from innate immune sensing. Cell 156, 134–145 (2014).

13. Crow, Y. J. & Stetson, D. B. The type I interferonopathies: 10 years on. Nat. Rev. Immunol. 22, 471–483 (2022).

14. Zhu, H. & Zheng, C. When PARPs Meet Antiviral Innate Immunity. Trends Microbiol. 29, 776–778 (2021).

15. Li, X. et al. cGAS guards against chromosome end-to-end fusions during mitosis and facilitates replicative senescence. Protein Cell 13, 47–64 (2022).

16. Song, J.-X., Villagomes, D., Zhao, H. & Zhu, M. cGAS in nucleus: The link between immune response and DNA damage repair. Front. Immunol. 13, 1076784 (2022).

17. Chen, H. et al. cGAS suppresses genomic instability as a decelerator of replication forks. Sci. Adv. 6, eabb8941 (2020).

18. Jiang, H. et al. Chromatin-bound cGAS is an inhibitor of DNA repair and hence accelerates genome destabilization and cell death. EMBO J. 38, e102718 (2019).

19. Humphries, F. & Fitzgerald, K. A. Loosening the grip on nuclear cGAS. Nat. Genet. 52, 1269–1270 (2020).

20. Dvorkin, S., Cambier, S., Volkman, H. E. & Stetson, D. B. New frontiers in the cGAS-STING intracellular DNA-sensing pathway. Immunity 57, 718–730 (2024).

21. Volkman, H. E., Cambier, S., Gray, E. E. & Stetson, D. B. Tight nuclear tethering of cGAS is essential for preventing autoreactivity. eLife 8, e47491 (2019).

22. Zierhut, C. et al. The Cytoplasmic DNA Sensor cGAS Promotes Mitotic Cell Death. Cell 178, 302–315.e23 (2019).

23. Boyer, J. A. et al. Structural basis of nucleosome-dependent cGAS inhibition. Science 370, 450–454 (2020).

24. Cao, D., Han, X., Fan, X., Xu, R.-M. & Zhang, X. Structural basis for nucleosome-mediated inhibition of cGAS activity. Cell Res. 30, 1088–1097 (2020).

25. Kujirai, T. et al. Structural basis for the inhibition of cGAS by nucleosomes. Science 370, 455–458 (2020).

26. Michalski, S. et al. Structural basis for sequestration and autoinhibition of cGAS by chromatin. Nature 587, 678–682 (2020).

27. Pathare, G. R. et al. Structural mechanism of cGAS inhibition by the nucleosome. Nature 587, 668–672 (2020).

28. Zhao, B. et al. The molecular basis of tight nuclear tethering and inactivation of cGAS. Nature 587, 673–677 (2020).

29. Wischnewski, M. & Ablasser, A. Interplay of cGAS with chromatin. Trends Biochem. Sci. 46, 822–831 (2021).

30. Nakatani, Y. & Ogryzko, V. Immunoaffinity purification of mammalian protein complexes. Methods Enzymol. 370, 430–444 (2003).

31. Jahan, C., et al. Unintegrated HIV-1 DNA recruits cGAS via its histone-binding domain to escape innate immunity. Proc. Natl. Acad. Sci. 122, e2424465122 (2025).

32. Liu, H. et al. Nuclear cGAS suppresses DNA repair and promotes tumorigenesis. Nature 563, 131–136 (2018).

33. Oberdoerffer, P. & Miller, K. M. Histone H2A variants: Diversifying chromatin to ensure genome integrity. Semin. Cell Dev. Biol. 135, 59–72 (2023).

34. Sun, Z. & Bernstein, E. Histone variant macroH2A: from chromatin deposition to molecular function. Essays Biochem. 63, 59–74 (2019).

35. Chen, H. et al. MacroH2A1.1 and PARP-1 cooperate to regulate transcription by promoting CBP-mediated H2B acetylation. Nat. Struct. Mol. Biol. 21, 981–989 (2014).

36. Gentili, M. et al. The N-Terminal Domain of cGAS Determines Preferential Association with Centromeric DNA and Innate Immune Activation in the Nucleus. Cell Rep. 26, 2377–2393.e13 (2019).

37. Chakravarthy, S. et al. Structural characterization of the histone variant macroH2A. Mol. Cell. Biol. 25, 7616–7624 (2005).

38. Chakravarthy, S. & Luger, K. The histone variant macro-H2A preferentially forms ‘hybrid nucleosomes’. J. Biol. Chem. 281, 25522–25531 (2006).

39. Stark, H. GraFix: stabilization of fragile macromolecular complexes for single particle cryo-EM. Methods Enzymol. 481, 109–126 (2010).

40. Huang, H. & Wu, Q. Pushing the TAD boundary: Decoding insulator codes of clustered CTCF sites in 3D genomes. BioEssays News Rev. Mol. Cell. Dev. Biol. 46, e2400121 (2024).

41. Hooftman, A., Keller, A. & Ablasser, A. The cGAS-STING pathway: Mechanism and medical implications. Cell 189, 3849–3870 (2026).

42. Li, H., Zhang, Y., Gao, S. & Yin, H. T cell-intrinsic cGAS-CTCF regulation maintains regulatory T cell development and function. Cell Rep. 44, 116302 (2025).

43. Tomaskovic, I. et al. DNA-protein cross-links promote cGAS-STING-driven premature aging and embryonic lethality. Science 391, eadx9445 (2026).

44. Sebastian, R. et al. Epigenetic Regulation of DNA Repair Pathway Choice by MacroH2A1 Splice Variants Ensures Genome Stability. Mol. Cell 79, 836–845.e7 (2020).

45. Ruiz, P. D., Hamilton, G. A., Park, J. W. & Gamble, M. J. MacroH2A1 Regulation of Poly(ADP-Ribose) Synthesis and Stability Prevents Necrosis and Promotes DNA Repair. Mol. Cell. Biol. 40, e00230–19 (2019).

46. Kim, J., Oberdoerffer, P. & Khurana, S. The histone variant macroH2A1 is a splicing-modulated caretaker of genome integrity and tumor growth. Mol. Cell. Oncol. 5, e1441629 (2018).

47. Kim, J. et al. The macroH2A1.2 histone variant links ATRX loss to alternative telomere lengthening. Nat. Struct. Mol. Biol. 26, 213–219 (2019).

48. Xu, C., Xu, Y., Gursoy-Yuzugullu, O. & Price, B. D. The histone variant macroH2A1.1 is recruited to DSBs through a mechanism involving PARP1. FEBS Lett. 586, 3920–3925 (2012).

49. Yang, H., Wang, H., Ren, J., Chen, Q. & Chen, Z. J. cGAS is essential for cellular senescence. Proc. Natl. Acad. Sci. U. S. A. 114, E4612–E4620 (2017).

50. Glück, S. et al. Innate immune sensing of cytosolic chromatin fragments through cGAS promotes senescence. Nat. Cell Biol. 19, 1061–1070 (2017).

51. Chen, Y.-A. et al. Extrachromosomal telomere repeat DNA is linked to ALT development via cGAS-STING DNA sensing pathway. Nat. Struct. Mol. Biol. 24, 1124–1131 (2017).

52. Chen, H. et al. MacroH2A1 and ATM Play Opposing Roles in Paracrine Senescence and the Senescence-Associated Secretory Phenotype. Mol. Cell 59, 719–731 (2015).

53. Ebata, H., Loo, T. M. & Takahashi, A. Telomere Maintenance and the cGAS-STING Pathway in Cancer. Cells 11, 1958 (2022).

54. Wang, X., Brea, L. T. & Yu, J. Immune modulatory functions of EZH2 in the tumor microenvironment: implications in cancer immunotherapy. Am. J. Clin. Exp. Urol. 7, 85–91 (2019).

55. Wang, Y., Bui, T. & Zhang, Y. The pleiotropic roles of EZH2 in T-cell immunity and immunotherapy. Int. J. Hematol. 116, 837–845 (2022).

56. Wu, B., Zhang, B., Li, B., Wu, H. & Jiang, M. Cold and hot tumors: from molecular mechanisms to targeted therapy. Signal Transduct. Target. Ther. 9, 274 (2024).

57. Pang, L. et al. Epigenetic regulation of tumor immunity. J. Clin. Invest. 134, e178540 (2024).

58. Fischetti, I., De Luca, M. & Jachetti, E. Exploring the role of EZH2 modulation in shaping the tumor microenvironment. Epigenomics 16, 1265–1268 (2024).

59. Zhou, L., Mudianto, T., Ma, X., Riley, R. & Uppaluri, R. Targeting EZH2 Enhances Antigen Presentation, Antitumor Immunity, and Circumvents Anti-PD-1 Resistance in Head and Neck Cancer. Clin. Cancer Res. Off. J. Am. Assoc. Cancer Res. 26, 290–300 (2020).

60. Gao, M. et al. Exploring the therapeutic potential of targeting polycomb repressive complex 2 in lung cancer. Front. Oncol. 13, 1216289 (2023).

