## Supplemental for "MacroH2A1 and H3K27me3 target cGAS to facultative heterochromatin to restrain innate immune activation"

**The PDF file includes:**

Materials and Methods

Figs. S1 to S7

References 61-77

**Other Supplementary Materials for this manuscript include the following:**

Table S1

Materials and Methods

**Plasmids.** Plasmids encoding cGAS were provided by N. Laguette. The cGAS gene was cloned into the pOZ-Puro-c-ter vector^30^ to generate a cGAS-Flag-HA expression plasmid. The mH2A1.1 plasmid was provided byA. G. Ladurner^61^. Subsequently, mH2A1.1 was cloned into the pOZ-IL2R-N-ter vector to generate an HA-mH2A1.1 expression plasmid.

**Cell lines**. 293T, MRC5, HeLa, and HeLa cGAS-KO cells reconstituted with either cGAS-Flag-HA or an empty vector control^31^ were cultured in Dulbecco’s modified Eagle medium (DMEM) supplemented with 10% FBS, 1 mM sodium pyruvate, and antibiotics. The Spodoptera frugiperda Sf9 cells and Trichoplusia ni High Five cells were cultured in SF-900III (Thermo Fisher) or ESF 921 (Expression Systems) medium, both containing 1× Penicillin-Streptomycin (Nacalai Tesque).

**Antibodies**: Antibodies used in this study: Western blot: anti-laminB1 (abcam, ab16048), anti-HA (BioLegend, 901503; Roche, 11867423001), anti-cGAS (Cell Signaling Technology [CST], 15102; Proteintech, 26416-1-AP), anti-mH2A1 (Abcam, ab37264), anti-mH2A1.1 (CST, 12455), anti-mH2A1.2 (CST, 9733), anti-H3K27me3 (CST, 9733), anti-H3K9me3 (Abcam, ab8898), anti-H3K9me2/3 (CST, 5327), anti-H3 (Abcam, ab1791), anti-H4 (Abcam, ab31830), anti-PARP1 (Santa Cruz Biotechnology [SCBT], sc-7150), anti-H3K36me3 (abcam, ab9050), anti-POLE3 (Bethyl, A301-245A), anti-HuR (sc-5261), anti-H2AX (abcam, ab11175), anti-γH2AX (millipore, 05-636), anti-IRF3 (CST, 4302), anti-phospho-IRF3 S386 (CST, 37829), anti-STING (CST, 13647), anti-phospho-STING S366 (CST, 19781). Endogenous/re-IP experiments: anti-HA (BioLegend, 901503), anti-H3K27me3 (CST, 9733), anti-H3K9me3 (Abcam, ab8898), anti-H3K36me3 (CST, 4909), anti-H3K4me3 (abcam, ab8580) and non-specific IgG (in-house). Beads: FLAG-M2 agarose beads (Sigma, F2426 or A2220) and HA agarose beads (SCBT, sc-7392AC) were used for FLAG and/or HA immunoprecipitation (IP).

**High-salt chromatin extraction (ChE).** All steps were performed on ice or in a cold room except for Nuclease S7 digestion. Immediately before use, the following additives were included in cold solutions: 5 mM β-glycerophosphate, 5 mM sodium butyrate, 5 mM β-mercaptoethanol, and 0.5 mM phenylmethylsulfonyl fluoride (PMSF). Cells were washed with cold PBS and resuspended in 5 pellet volumes of Extraction Buffer 1 (PBS, 0.2 mM EDTA, 0.1% Triton X-100), followed by incubation for 10 minutes on a rotating wheel. After centrifugation (1,000 × g, 10 minutes), the pellet was extracted with 4 pellet volumes of Extraction Buffer 2 (20 mM Tris [pH 7.4], 25% glycerol, 1.5 mM MgCl₂, 0.2 mM EDTA, 400 mM KCl) for 30 minutes on a rotating wheel. Following centrifugation (1,000 × g, 10 minutes), the pellet was treated with 6 pellet volumes of Extraction Buffer 3 (3 mM EGTA, 0.2 mM EDTA)^21^ for 30 minutes on a rotating wheel. The pellet was then resuspended in 3 volumes of Extraction Buffer 4 (50 mM Tris [pH 8], 0.05% IGEPAL 630, 1 mM MgCl₂, 5 mM CaCl₂, 2 U/µL Nuclease S7 [Roche, #10107921001]) and incubated sequentially: 1.5 minutes at 37°C, 20 minutes at room temperature (RT), and 40 minutes at 4°C. After centrifugation (1,000 × g, 10 minutes), the pellet was extracted with 3 volumes of Extraction Buffer 5 (50 mM Tris [pH 8], 0.05% IGEPAL 630, 500 mM NaCl) for 30 minutes on a rotating wheel. The supernatant was retained on ice, and the pellet was washed with the same volume of Extraction Buffer 6 (20 mM Tris [pH 7.4], 25% glycerol, 1.5 mM MgCl₂, 20 mM KCl, 0.2 mM EDTA, 2 mg/mL BSA, 0.2% Triton X-100). After centrifugation (1,000 × g, 10 minutes), the final supernatant was combined with the previous supernatant and centrifuged at maximum speed for 30 minutes. The resulting chromatin extract (ChE) was flash-frozen in liquid nitrogen.

**Tandem affinity chromatography**. Tandem affinity chromatography was performed as previously described^30^ using a 1:100 bead-to-ChE ratio. Ten percent of the FLAG- and HA-immunoaffinity-purified cGAS-Flag-HA or mock immunoprecipitates (derived from ten 15 cm plates) were resolved by SDS-PAGE and stained with the SilverQuest™ Silver Staining Kit (Invitrogen). The remaining eluate was stained with Colloidal Blue™ Stain (Invitrogen). Four colloidal blue-stained gel regions were excised and analyzed by tandem mass spectrometry at the Taplin Biological Mass Spectrometry Facility, Harvard Medical School (Boston, MA, USA).

**Reciprocal immunoprecipitation**. The H3K9me3 antibody was crosslinked to protein A Sepharose beads (Sigma, #P9424) using a dimethyl pimelimidate (DMP; Sigma). For reciprocal immunoprecipitation, 40 μl of FLAG-purified cGAS-Flag-HA complex was diluted in 400 μl Buffer B (20 mM Tris-HCl [pH 7.5], 10% glycerol, 150 mM KCl, 5 mM MgCl₂, 0.5 mM EDTA, 0.1% Tween 20, 0.05% Triton X-100, 5 mM β-mercaptoethanol, 0.5 mM PMSF) containing 0.1 mg/ml BSA and incubated with the indicated antibodies under gentle agitation at 4°C. After washing with Buffer B, the beads were boiled in Laemmli loading buffer.

**Immunoprecipitation of endogenous proteins**. Extracts were prepared as described above, except 50 mM iodoacetamide (Sigma, #I6125) was added prior to EDTA/EGTA treatment, and lysis was performed in PBS containing 400 mM NaCl (instead of Extraction Buffer 2). A total of 100 μL of ChE was incubated with the primary antibody at 4°C for 4 hours. During antibody incubation, protein A beads were saturated in PBSTT300 buffer (PBS with 300 mM NaCl, 0.1% Tween 20, 0.05% Triton X-100) containing 0.1 mg/ml BSA and 0.1% polyvinylpyrrolidone (PVP40; Sigma) on a wheel at 4°C. After 4 hours, the saturation buffer was discarded. The ChE/antibody mixture was diluted in 500 μL of PBSTT300 supplemented with 0.1 mg/ml BSA, combined with saturated beads, and incubated on a wheel at 4°C for 45 minutes. Beads were washed four times in PBSTT300 and boiled in Laemmli loading buffer.

**Protein purification and heterotypic nucleosome reconstitution**. Streptavidin-tagged human histone H2A, mH2A1.1, mH2A1.2 and streptavidin-tagged H2B were subcloned into pAC8-derived vectors respectively (pAC8/strep-H2A, pAC8/strep-mH2A1.1, pAC8/strep-mH2A1.2, pAC8/strep-H2B). *Spodoptera frugiperda* Sf9 cells were co-transfected with pAC8 and viral DNA, and then incubated at 27°C for 5 days. The supernatant (P0 virus) of the medium, which contains the baculovirus expressing strep-H2A, strep-mH2A1.1, strep-mH2A1.2 or strep-H2B, was collected. The virus was successively amplified by infecting and incubating Sf9 cells for 3 days. Such procedures were repeated until the P3 virus was obtained. The baculovirus of strep-H2A or strep-mH2A1.1 or strep-mH2A1.2 and the baculovirus of strep-H2B were co-expressed in 400 mL cultures of *Trichoplusia ni* High Five cells, using the Bac-to-Bac system (Thermo Fisher). After incubating for 3 days at 27°C, the cells were lysed with lysis buffer (50 mM Tris-HCl [pH 8.0], 500 mM NaCl) containing 1 × protease inhibitor cocktails (Sigma), centrifuged, and the supernatant was loaded onto a Streptactin (IBA) affinity column. After extensive wash with Lysis buffer, H2A-H2B, mH2A1.1-H2B, and mH2A1.2-H2B were eluted with 2.5 mM desthiobiotin (IBA) in Lysis buffer containing 250 µM TCEP. The elutions were further purified on a size exclusion column (HiLoad 16/600 Superdex 200; Cytiva) to fractionate H2A-H2B, mH2A1.1-H2B or mH2A1.2-H2B dimers.

Histone H3.1 and H4 were purified from *E. coli* by the method previously described^62^. Equimolar amounts of lyophilized H3.1 and H4 were mixed in denaturing buffer (20 mM Tris-HCl (pH 7.5), containing 7 M guanidine hydrochloride and 20 mM 2-mercaptoethanol). After dialysis against refolding buffer (2 M NaCl, 10 mM Tris-HCl [pH 7.5], 1 mM EDTA, 2 mM 2-mercaptoethanol), H3.1-H4 tetramer was purified by size exclusion column chromatography (HiLoad 16/600 Superdex 200; Cytiva).

H2A-H2B dimers, mH2A1.1-H2B dimers and H3.1-H4 tetramer were mixed at 1:1 molar ratio and incubated on ice for 5 minutes to assemble heterotypic histone octamer. The histone octamers were then combined with 193-bp 601DNA, and nucleosomes containing mH2A1.1 were reconstituted by salt dialysis. The reconstituted nucleosomes were purified by 5% non-denaturing polyacrylamide gel electrophoresis, using a Prep Cell apparatus (Bio-Rad) as previously described^63^. Nucleosomes containing mH2A1.2 was reconstituted using the same procedure.

Recombinant cGAS lacking unstructured N-terminal region protein was purified as the same method described previously^25^.

**Gel electrophoretic mobility shift assays (EMSAs)**. Various concentrations of cGAS (0, 100, 200, 300, or 400 nM) were mixed with, mH2A1.1-nucleosomes (200 nM), or mH2A1.2-nucleosomes (200 nM), or canonical nucleosomes (200 nM) in binding buffer (60 mM NaCl, 20 mM Tris-HCl, pH 7.5, 0.2 mM TCEP, 0.2 mM DTT, 4% glycerol) and incubated for 35 minutes at 37°C. The mixtures were fractionated on a 4% non-denaturing polyacrylamide gel in 0.5 × TBE buffer (44.5 mM Tris, 44.5 mM boric acid, 1 mM EDTA). The gel was stained with Ethidium Bromide and visualized with a Typhoon imaging analyzer (Cytiva). The intensities of interested bands were quantified by Image Quant software and Graph representations were computed by Prism10 software.

**Cryo-EM sample preparation**. For the structure determination of the cGAS-mH2A1.1Nuc complex, cGAS and mH2A1.1nucleosomes were mixed and the cGAS-mH2A1.1Nuc complex was isolated by GraFix method described previously^25^. Fractions containing the cGAS-mH2A1.1Nuc complex were combined and concentrated. 2.5 µL of the eluate sample was applied to R 1.2/1.3 Quantifoil grids (Quantifoil Micro Tools GmbH). Glow discharging was performed with a PIB-10 ION Bombarder (Vacuum Device) for 90 seconds under hard conditions. After 7 seconds of blotting, the grid was vitrified using a Vitrobot Mark IV (Thermo Fisher) at 4°C and 100% humidity.

**Cryo-EM data acquisition and image processing**. Micrographs were captured by a Krios G4 transmission electron microscope (Thermo Fisher) with a K3 direct electron detector and a BioQuantum energy filter (Gatan). 21,681 movies were captured for each cryo-EM sample by the EPU software (Thermo Fisher). Data acquisition was performed with a 1.06 Å pixel size and -1.0 to -2.5 µm defocus values at 0.25 µm steps. Data collection was performed by 40 frames. CryoSPARC was used for further data processing^64^. Motion correction and CTF estimation were performed using Patch Motion Correction^65^ and Patch CTF Estimation^66^, respectively. The cGAS-mH2A1.1Nuc complex particles were picked by Blob Picker using a Laplacian of Gaussian filter. The particles were extracted with a box size of 320, then decreased the box size to 80 by fourier cropping and performed 2D classification. After extensive removal of non-cGAS-mH2A1.1Nuc complex particles, heterogeneous refinement was performed. The cGAS-mH2A1.1Nuc complex class was selected and performed homogeneous refinement. Particles were further re-extracted with a box size of 400 and performed several rounds of 3D classification with a mask. The cGAS-mH2A1.1Nuc complex class was selected and performed Non-uniform Refinement. Then Reference Based Motion Correction, Local Motion Correction were applied. Final particle sets were analyzed by Non-uniform Refinement and the DeepEMhancer software^67^.

**Model building and refinement**. The initial model of cGAS-mH2A1.1Nuc was built based on the structures previously determined (PDB IDs: 7C0M, 9J8M and 2F8N). Models of amino acids were built by real space refinement with the Phenix software^68^, and COOT^69^. Structural figures were displayed by the Pymol^70^ and ChimeraX software^71^.

**cGAS-chromatin extraction and Glycerol gradient**. cGAS-Chromatin Extraction: Salt extraction was performed as described^21^. Briefly, cell pellets were washed in PBS, resuspended in cytoplasmic extraction buffer (10 mM HEPES [pH 7.9], 10 mM KCl, 1.5 mM MgCl₂, 0.34 M sucrose, 10% glycerol, 0.2% NP-40, protease inhibitors), and incubated for 10 minutes on ice with occasional vortexing. Nuclei were centrifuged at 6,500 × g for 5 minutes at 4°C, and the cytoplasmic fraction was collected. Nuclei were washed in cytoplasmic extraction buffer without NP-40 and centrifuged again at 6,500 × g for 5 minutes at 4°C. Nuclei were then resuspended in salt extraction buffer (3 mM EDTA, 0.2 mM EGTA, protease inhibitors), vortexed for 1 minutes (10secondson, 10secondsoff), and incubated on ice for 30 minutes with vortexing every 10 minutes for 15 sec. Lysates were centrifuged at 6,500 × g for 5 minutes at 4°C, and the no-salt fraction was collected. Pellets were successively resuspended in salt buffer (50 mM Tris-HCl [pH 8.0], 0.05% NP-40) containing increasing NaCl concentrations (0.25, 0.5, 0.75, and 1 M). For each concentration, pellets were incubated on ice for 15 minutes (vortexed every 5 minutes for 15 sec), then centrifuged at maximum speed for 5 minutes at 4°C. Supernatants were collected for each NaCl concentration. The final pellet was resuspended in salt buffer containing 1 M NaCl, sonicated using a Bioruptor Pico (Diagenode; 30secondson/30secondsoff, 4 cycles), and centrifuged at 16,000 × g for 20 minutes to collect the supernatant. All fractions were analyzed by Western blot using anti-cGAS (15102, Cell Signaling Technology) and anti-H3 (4620, CST).

Glycerol Gradient: A total of 200 μL of FLAG-purified material was loaded onto a 4.5 mL 10–40% glycerol gradient (40%: 0.5 mL; 35%: 0.5 mL; 30%: 0.5 mL; 25%: 1 mL; 20%: 1 mL; 10%: 1 mL) in buffer (20 mM Tris [pH 7.5], 150 mM NaCl, 5 mM MgCl₂, 0.1% Tween 20) and centrifuged at 55,000 rpm for 4 hours at 4°C in a Beckman SW55Ti rotor. Fractions (200 μL each) were collected from the gradient top. Proteins were concentrated via TCA precipitation: samples were treated with 5% TCA, incubated on ice for 30 minutes, and centrifuged at 16,000 × g for 20 minutes at 4°C. Pellets were washed twice with –20°C acetone (centrifugation: 16,000 × g, 5 minutes, 4°C), dried briefly at 95°C, and resuspended in 20 μL PBS. Equal volumes of each fraction were analyzed by Western blot using specific antibodies.

**IFN-related genes (ISG) quantification**. RNA was purified from mH2A1 siRNA-transfected cells or UNC1999-treated cells. HeLa cells were transfected with the following siRNAs for 72 hours: Non-targeting (NT) control (Dharmacon, #D-001810-01), mH2A1.1 #1: GUUGUACAGGCUGACAUUG-dTdT^72^, mH2A1.1 #2: GUUGUACAGGCUGACAUUG-dTdT^72,73^, mH2A1.1 #3: CAUUGCCUCGAUCGACAGU-dTdT, mH2A1.2: GCUUUGAGGUGGAGGCCAU-dTdT^47^. In experiments involving cGAS or STING inhibitors (Invivogen), cells were pretreated with 10 µM G140 and 2 µM H151 for 3 hours before siRNA transfection; inhibitors were maintained throughout the experiment. HeLa cells were treated with 5 µM UNC1999 (Sigma-Aldrich) for 48 hours. RNA was purified using the RNeasy Plus Mini Kit (Qiagen) according to the manufacturer’s instructions. cDNA was synthesized using SuperScript™ IV Reverse Transcriptase (Thermo Fisher Scientific). The mRNA expression of ISGs and GAPDH was quantified by qPCR (Roche LightCycler®) with the following primers: ISG15 (Forward: CAGCCATGGGCTGGGAC. Reverse: GCCGATCTTCTGGGTGATCT). IFIT1 (Forward: GGAATACACAACCTACTAGCC. Reverse: CCAGGTCACCAGACTCCTCA. MXA (Forward: ATGAGCTAATCACCCTGGAG. Reverse: ATACCCAATGTCAGCAGGC). GAPDH (Forward: CTGGCGTCTTCACCACCATGG. Reverse: CATCACGCCACAGTTTCCCGG)^11^. IFIT2 (Forward: GGAGCAGATTCTGAGGCTTTGC. Reverse: GGATGAGGCTTCCAGACTCCAA). MX2 (Forward: CAGAGGCAGCGGAATCGTAA. Reverse: TGAAGCTCTAGCTCGGTGTTC). RSAD2 (Forward: GCTCTAAGAGAAGCAGAAAG. Reverse: CATCTTCTGGTTAGATTCAGG). IFNβ (Forward: AAACTCATGAGCAGTCTGCA. Reverse: AGGAGATCTTCAGTTTCGGAGG).

**Chromatin Immunoprecipitation (ChIP) assay**. ChIP assays were carried out on HeLa KO cGAS and HeLa-cGAS-Flag-HA transfected with NT, mH2A1.1 or mH2A1.2 siRNA for 48 hrs. ChIP for histone H3, histones modifications H3K37me3 and H3K9me3, mH2A1.1 and mH2A1.2 were performed using SimpleChIP Plus Enzymatic Chromatin IP Kit (Cell Signaling Technology) according to manufacturer protocol^74^. Briefly, 10 µg of MNase digested chromatin were reacted with anti-H3 (4620, Cell Signaling Technology), anti-H3K27me3 (9733, Cell Signaling Technology), anti-H3K9me3 (ab8898, abcam), anti-HA (ab9110, abcam), mH2A1.1 (12455, Cell Signaling Technology) or mH2A1.2 (ab37264, abcam) antibodies. qPCR was performed using specific primers : α-satellite (α-satellite-F : AGGGAATGTCTTCCCATAAAAACT, α-satellite-R : GTCTACCTTTTATTTGAATTCCCG)^75^; GAPDH-pro (GAPDH-pro-F : TACTAGCGGTTTTACGGGCG, GAPDH-pro-R : TCGAACAGGAGGAGCAGAGAGCGA) (Sigma); MYT-1 (MYT-1-F : AGGCACCTTCTGTTGGCCGA, MYT-1-R : AGGCAGCTGCCTCCCGTACA); b13 (b13-F : GAAACCACCCACGTGTTCTTTC, b13-R : TGGATCCCTGATCACCTCCAAG)^74^ and IL8 (IL8-2kb-F : AGGCAACCGTTAGGGAAAAG, IL8-2kb-R : GGACACAACCTGGCTTGACT)^53^. Libraries were prepared and sequenced by Novogene (NovaSeq X Plus PE150).

**ChIPseq analyses**. All paired-end fastq files produced in this study were aligned to the GRCh38 genome assembly using bowtie2 (v2.4.2, PMID: 22388286) with the following parameters -q --very-fast -k 1. The resulting SAM files were converted to BAM files using samtools view function (v1.11, PMID: 33590861) with high quality parameters (-q 30 -F 4 -b). Peak calling was performed using MACS2 (v2.1.1, PMID: 18798982) with callpeak parameters -f BAMPE -q 0.01 --SPMR -B. Wig normalized coverage profiles produced by MACS2 were imported into R using the misha genomics framework (v4.0.6, https://tanaylab.github.io/misha/) and used for all downstream analyses and quantifications after selecting filtering MACS2 narrow peaks (parameters: qValue between 5 and 100). To further validate the specificity of the detected Flag-HA binding sites, a z-score was calculated between cGAS- Flag-HA ChIP signal obtained in WT cells as compared to the signal obtained following the same procedure in cGAS-KO cells. The z-scores were calculated for every peak (peak summit +/- 500bp) using the cGAS filtered peak. Only peaks with a z-score higher than 1 were maintained (Supplementary Fig. S5A). As a control, we also computed peaks on the cGAS-KO sample using the reciprocal approach (Supplementary Fig. S5A) giving raise to the ‘Control Peaks’ group. The ‘Random Peaks’ group refers to a randomized set of genomic coordinates computed to match the numbers and chromosome distribution of the filtered WT-cGAS peaks.

To assess for potential enrichment of cGAS enriched sites, we computed overlap percentages with a series of genome annotations such as TSS (UCSC canonical genes), Centromeres (UCSC), simple repeats (UCSC) and a comprehensive set of transcription factor ChIPseq peaks available for HeLa-S3 sets through the ENCODE project data portal. The distance threshold for a positive match in all cases was 10kb from the cGAS filtered peak summits and was based on the ginterval.neighbors function (misha). P-Values were computed as the fraction of equal or higher percentages obtained in 200 iterations using chromosome matched randomized genomic coordinates.

For peak clustering, we utilized the normalized ChIP signal across all 5kb regions centered on the filtered cGAS peak summits by extracting profiles at 50bp resolution for cGAS, mH2A1.1, mH2A1.2, H3K27me3 and H3K9me3 using in-house R scripts based on the misha gextract function. The resulting matrices for cGAS, H3K27me3 and H3K9me3 signals were merged into a single matrix that was clustered using R’s kmeans function with k=8 and set.seed.

Differential ChIP heatmap profiles were computed by extracting the normalized ChIP signal across all 5kb regions centered on the filtered cGAS peak summits at 50bp resolution using in-house R scripts based on the misha gextract function. The differential matrix profile was computed by subtracting the WT matrix from the KO matrix. Average per cluster variation was computed as -log2(KO/WT) across the average ChIP signal of 500bp regions centered on cGAS peak summit clusters using in-house R scripts based on the misha gextract function. Graph representations were computed using the iGraph package (v1.1.2, https://cran.r-project.org/web/packages/igraph/index.html).

**Publicly available datasets (TABLE - GEO)**. cryo-EM map and coordinate of the structure of cGAS-mH2A1.1Nuc was deposited in the Electron Microscopy Data Bank (EMDB) and Protein Data Bank (PDB) with the accession codes EMD-64189 (9UIA).

ChIPseq data from HeLa-S3 cells were obtained from the ENCODE Data Portal^76^. For all datasets, we used the latest version of the GRCh38 processed data including for signal (fold change over control, bigWig, combined replicates) and for peaks (bed narrowPeak). Default peaks were further filtered for scores between X and Y. All bigWig files were converted to misha tracks using the gtrack.import function.

ChIPseq profile for gammaH2AX (GSE60526) and for CENPA (GSE111381) were re-analyzed using the same pipeline used for the datasets produced in this study and converted to misha tracks using the gtrack.import function. All subsequent analyses were performed using in-house R scripts based on the misha gextract function.

**MicroC**. HeLa-S3 microC valid pairs were obtained from the 4D Nucleome Data Portal^77^. The interactions pairs were imported into the misha database using the gtrack.2d. import_contacts function. Expected profiles were computed using the shaman package (https://github.com/tanaylab/shaman). Average maps were computed at 1Mb regions centered on different cGAS peak summits subsets at a 10kb resolution. In all maps the -log2(Observed / Expected) ratio is represented. Genomic insulation profiles were computed on the Observed tracks using the gtrack.2d.gen_insu_track function of the shaman package with a 250kb window and 2kb resolution.

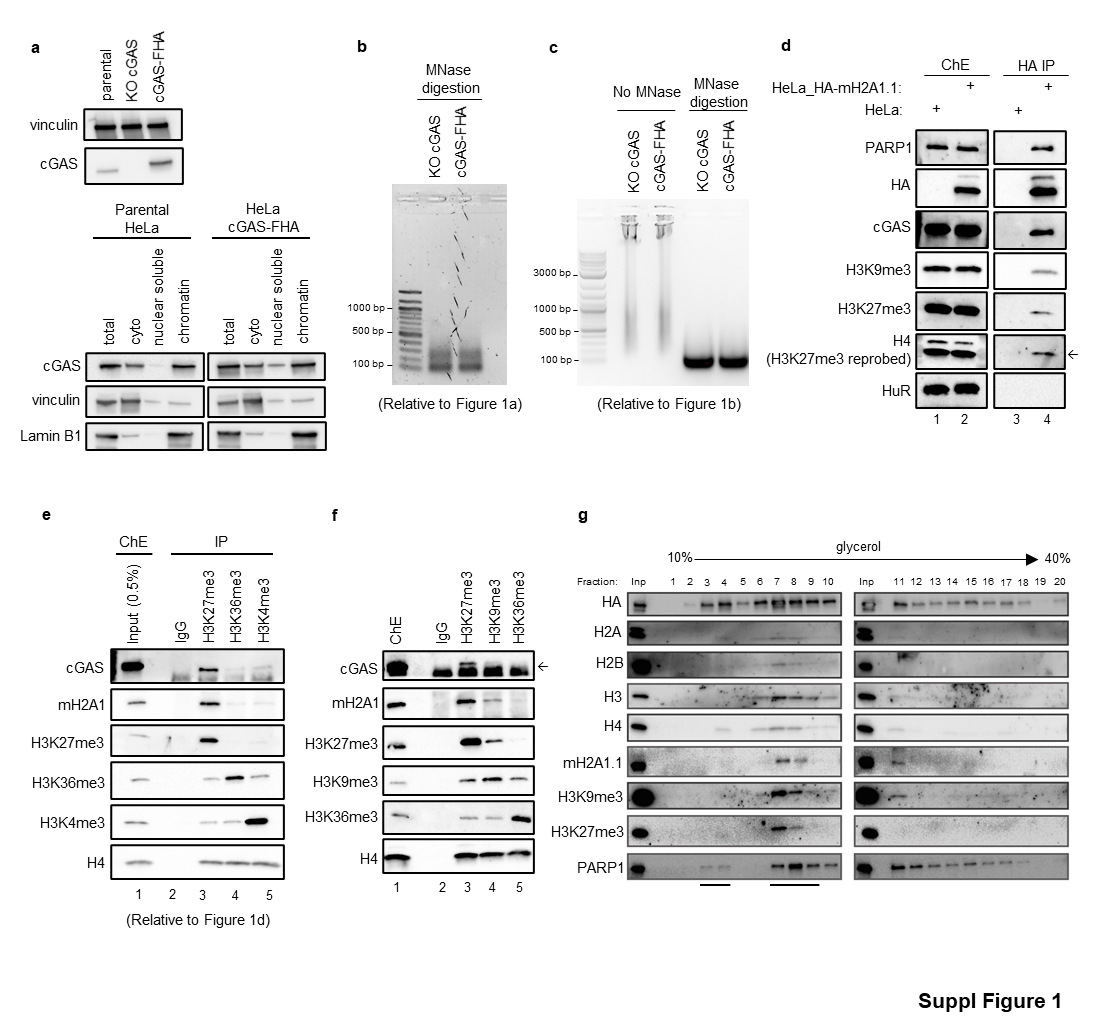

Fig. S1.

**a,** Characterization of HeLa cGAS-KO cells stably expressing cGAS-Flag-HA compared to parental HeLa cells. Immunoblot of cGAS expression (top panel). fractionation into cytosolic, nuclear soluble, and chromatin fractions (bottom panel). **(b, c)** Agarose gel showing the chromatin fragment size present in extracts used for cGAS-Flag-HA immunopurification (b, relative to Fig.1a) and extracts used for co-immunoprecipitation experiments (c, relative to Fig 1b). **d,** ChE of HeLa cells stably expressing HA-mH2A1.1 or untransduced control cells were subjected to HA IP. ChE (lanes 1-2) and HA-elutes (lanes 3-4) were resolved on SDS-PAGE and immunoprobed with the indicated antibodies. **e,** Endogenous immunoprecipitations on HeLa ChE using control rabbit IgG, or H3K27me3-, or H3K36me3-, or H3K4me3-specific antibodies, analyzed by immunoblotting with the indicated antibodies. **f,** Endogenous immunoprecipitations on MRC5 ChE extract using control rabbit IgG, H3K27me3-, H3K9me3-, or H3K36me3-specific antibodies was analyzed by immunoblotting using the indicated antibodies. **g,** Flag-purified cGAS-Flag-HA associated complexes were separated by centrifugation through a 10-40% glycerol gradient. Collected fractions were analyzed by Western blot using the indicated antibodies.

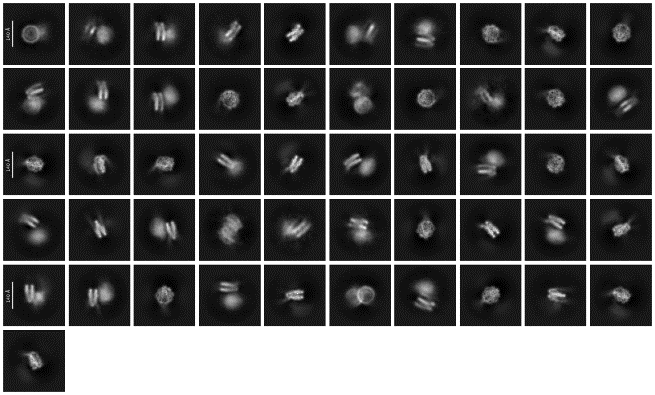

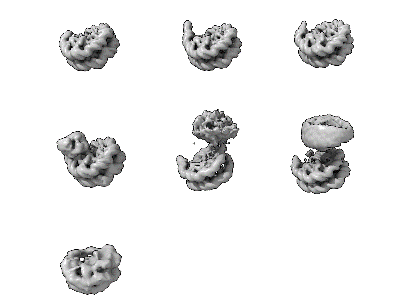

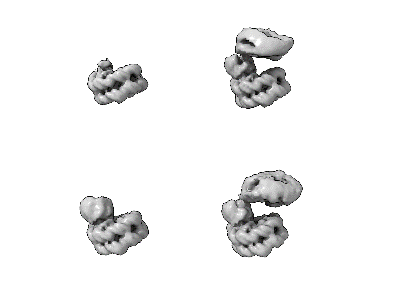

21,681 micrographs

Motion correction

CTF estimation

Blob picker

8.827,248 particles

2D classification

5,080,053 particles

Heterogeneous refinement (Fourier cropped)

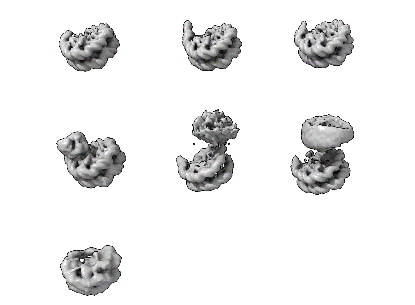

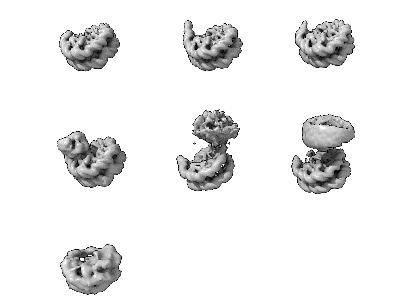

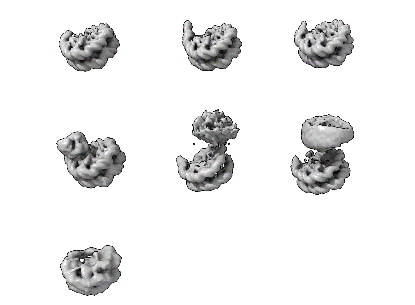

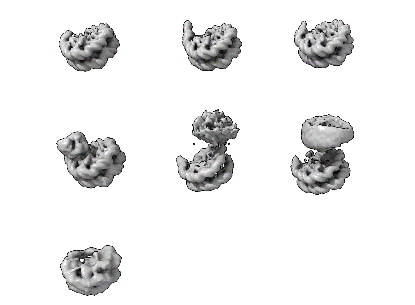

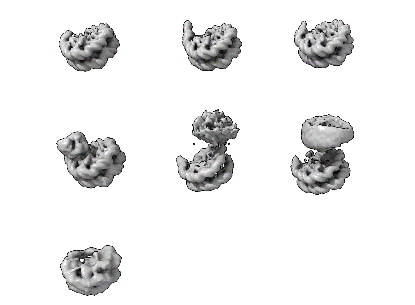

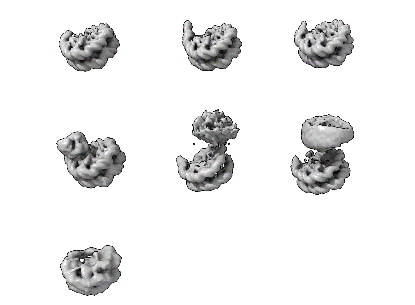

Class 0

Class 1

Class 2

Class 3

Class 4

Class 5

Class 6

3D classification (with mask)

Class 0

Class 1

Class 2

Class 3

18.9%

20.9%

29.9%

20.1%

5.5%

9.4%

4.1%

19.1%

18.9%

32.1%

21.1%

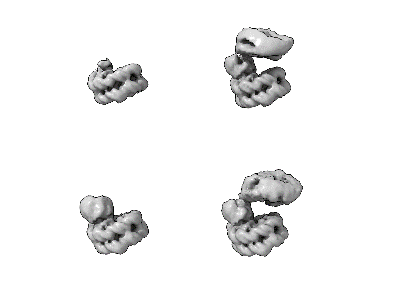

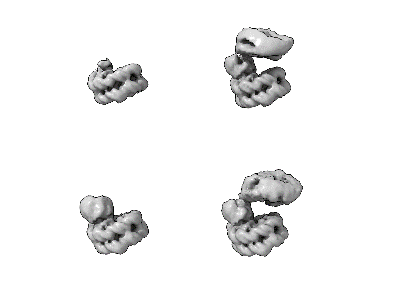

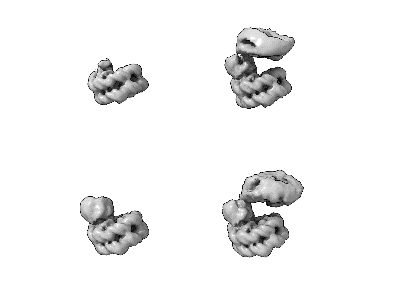

Reference Based Motion Correction

Local Motion Correction

Non-uniform Refinement

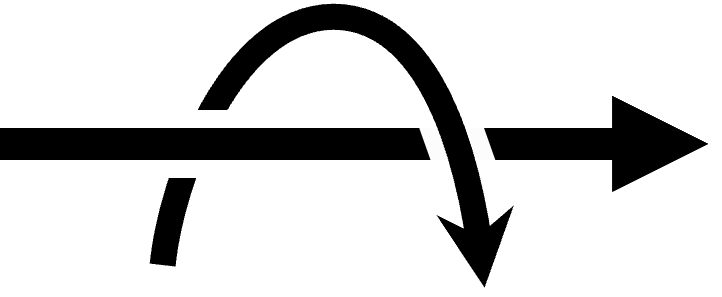

90°

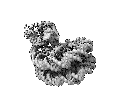

2.88 Å

316,743 particles

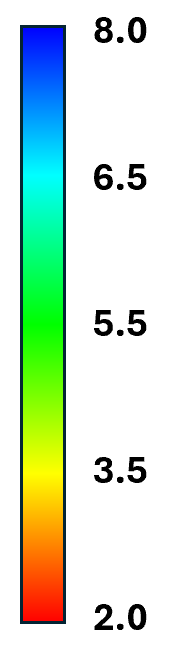

**a**

**b**

**c**

**d**

**e**

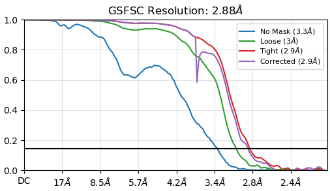

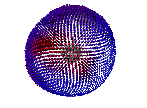

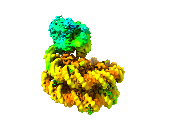

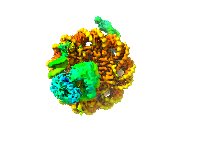

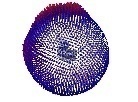

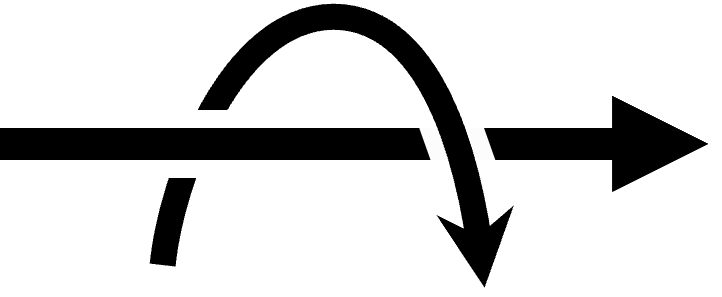

90°

**f**

(Å)

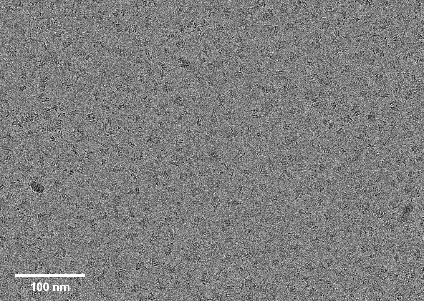

Fig. S2.

**a,** Entire cryo-EM procedures for structure determination of the cGAS-mH2A1.1Nuc complex by CryoSPARC. After motion correction and CTF estimation, particles were autopicked and extracted, and 2D classification was further performed. Selected particles were classified by Hetero Refinement and then refined by Homo Refine. 3D classification was performed and 7 classes were obtained. After one more 3D classification with mask, 4 classes were obtained. Selected class was processed by Reference Motion Correction and Local Motion Correction, and then refined by NU-Refine. The 2.88 Å resolution map was obtained as the final cryo-EM map and sharpened with deepEMhancer. **b,** Representative cryo-EM micrograph for the cGAS-mH2A1.1Nuc complex. **c,** 2D classification images after initial particle extraction. **d,** Gold-standard Fourier shell correlation curves (FSCs) for the cGAS-mH2A1.1Nuc complex. **e,** Local-resolution map of the cGAS-mH2A1.1Nuc complex. **f,** Angular distribution of the particles of the final map.

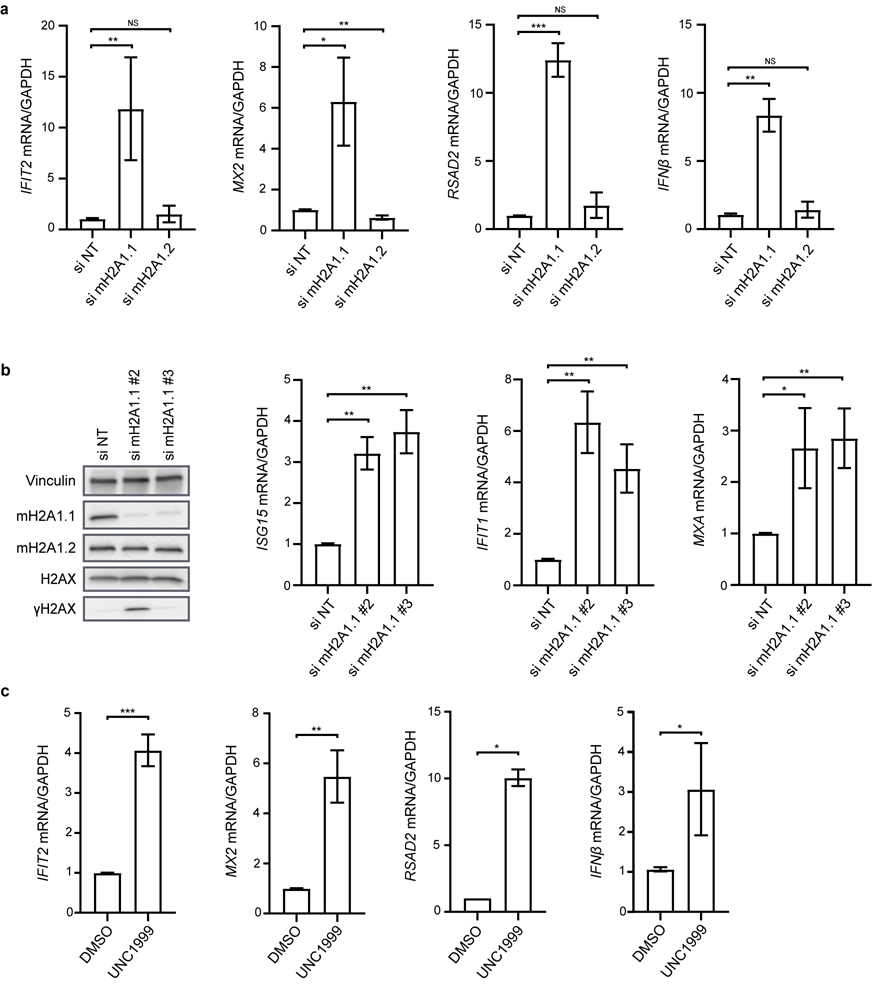

Fig. S3.

**a,** IFIT2, MX2, RSAD2 and IFNβ mRNA expression were measured by RT-qPCR (normalized to GAPDH mRNA). Mean ± SD of four independent experiments (relative to Fig. 4a). **b,** HeLa cells were transfected with two individual mH2A1.1 siRNA. ISG15, IFIT1, and MXA mRNA expression were measured by RT-qPCR (normalized to GAPDH mRNA). Mean ± SD of four independent experiments. mH2A1.1 depletion was assessed by Western blot analysis. One representative experiment of four independent experiments is shown. **c,** IFIT2, MX2, RSAD2 and IFNβ mRNA expression were measured by RT-qPCR (normalized to GAPDH mRNA). Mean ± SD of four independent experiments (relative to Fig. 4c).*P < 0.05, **P < 0.01, ***P < 0.001; independent Student’s t-test.

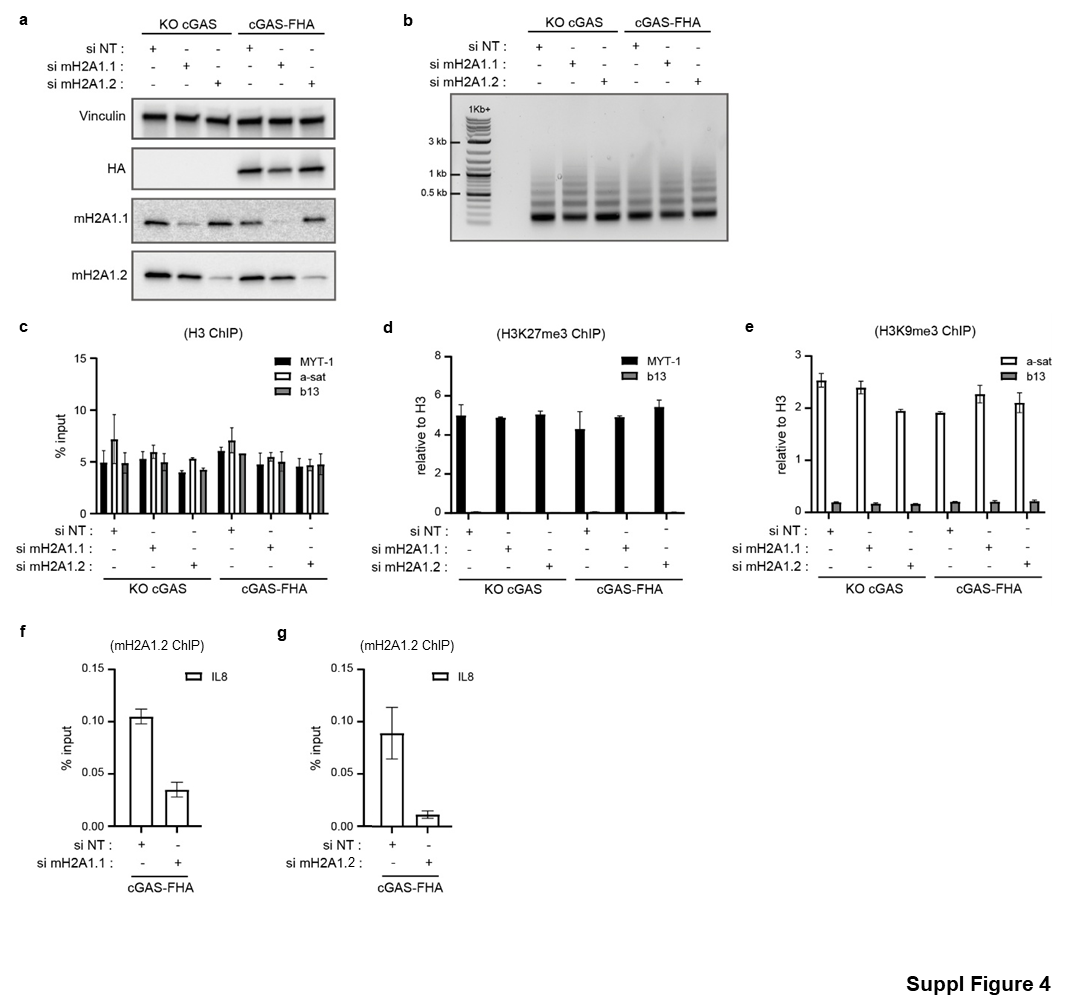

Fig. S4.

ChIP assay in cGAS-KO or cGAS-Flag-HA HeLa cells transfected with control (NT), mH2A1.1-, or mH2A1.2-specific siRNA for 48 h. **a,** Western blot analysis assessing expression of cGAS-Flag-HA and efficiency of mH2A1.1/mH2A1.2 depletion. One representative experiment of 2 is shown. **b,** MNase-digested chromatin profiles (used for ChIP and ChIP-seq experiments) were analyzed by agarose gel electrophoresis. One representative experiment of 2 is shown. **c–g,** ChIP assays were performed using anti-H3 (c), anti-H3K27me3 (d), anti-H3K9me3 (e), anti-mH2A1.1 (f), and anti-mH2A1.2 (g) antibodies and analyzed by qPCR with the indicated primers. Data represent mean ± SD of 2 independent experiments.

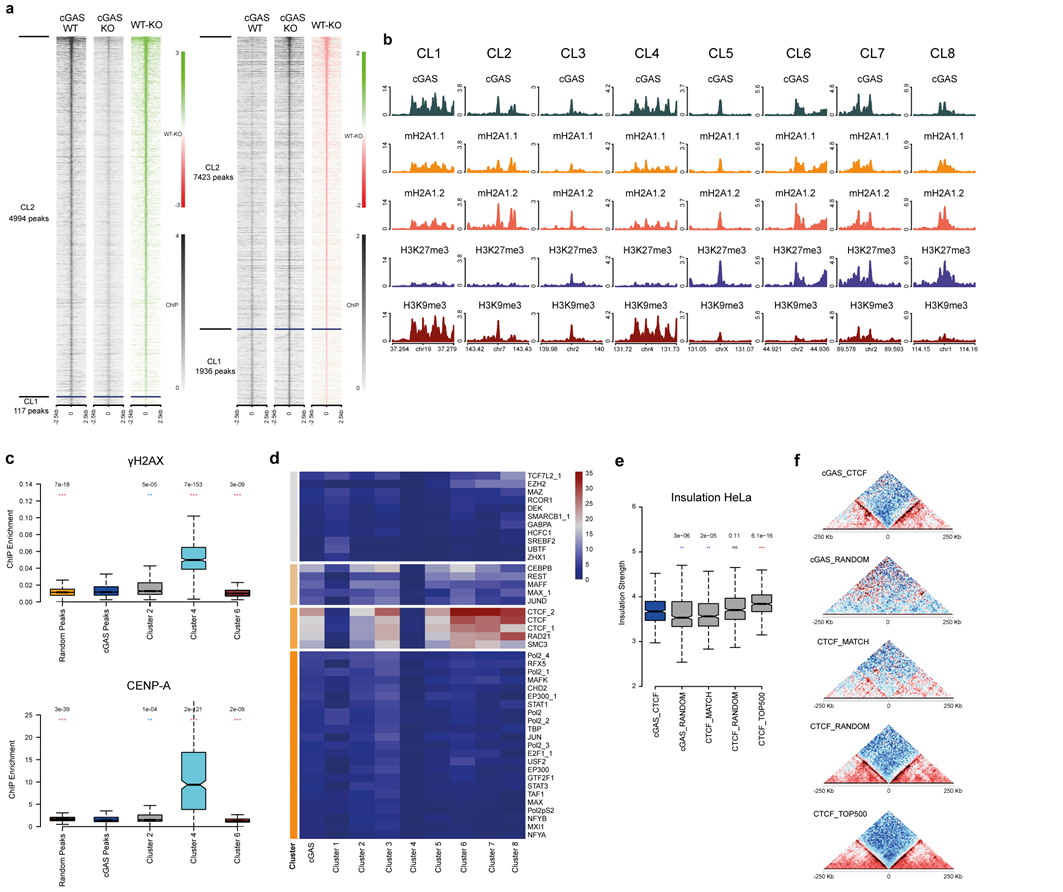

**Fig. S5.**

a, cGAS enrichment profiles across peaks identified in cGAS-Flag-HA (cGAS-WT, left panels) and cGAS-KO cells (right panels). Peaks are ranked by increasing (cGAS_WT) or decreasing (cGAS_KO) z-scores. Clusters are defined by a z-score cut-off of 1 (Z > 1 in cGAS_WT and Z < −1 in cGAS_KO). The resulting peaks are considered bona fide cGAS peaks (cGAS_WT) or cGAS control peaks (cGAS_KO) in all subsequent analyses. Differential enrichment profiles (WT – KO) are also illustrated for the same regions. b, Representative examples of cGAS peaks assigned to each of the 8 epigenetic clusters (Fig. 5c). Each vertical panel illustrates the cGAS, mH2A1.1, mH2A1.2, H3K27me3, and H3K9me3 enrichment profiles over a 15 kb genomic region centered on the peak summit. Clusters are ordered horizontally from left (Cluster 1) to right (Cluster 8). c, Boxplots comparing ChIP-seq signal distribution of γ-H2AX (top) and CENPA (bottom) across random genomic regions, cGAS peaks, and selected epigenetic clusters representing distinct epigenetic profiles (Cluster 2: H3K9me3; Cluster 4: centromeres; Cluster 6: H3K27me3). P-values were obtained by comparing the distribution of each cluster to all cGAS peaks (in blue) using a Wilcoxon two-sided test. d, Functional characterization of identified cGAS peaks and respective epigenetic clusters through k-means stratification of their intersection profiles with transcription factor (TF) peaks in HeLa cells (ENCODE IDR narrow peak datasets). Overlaps were computed using a 10 kb maximum distance between the cGAS peak summits and the center of the IDR narrow peaks. For simplicity, ENCODE peak datasets with fewer than 300 peaks or minimum overlaps (across all cGAS peaks or individual clusters) lower than 3% were excluded. e, Genomic insulation profiles across cGAS peaks overlapping CTCF sites, cGAS random peaks, CTCF peaks with matched (low) enrichment scores, random CTCF peaks, and top 500 scoring CTCF peaks. P-values were obtained by comparing the distribution of each cluster to cGAS peaks overlapping CTCF peaks (in blue) using a Wilcoxon two-sided test. f, Aggregate micro-C maps centered on cGAS peaks overlapping CTCF sites, cGAS random peaks, CTCF peaks with matched enrichment scores, random CTCF peaks, and top 500 scoring CTCF peaks (top to bottom panels, respectively).

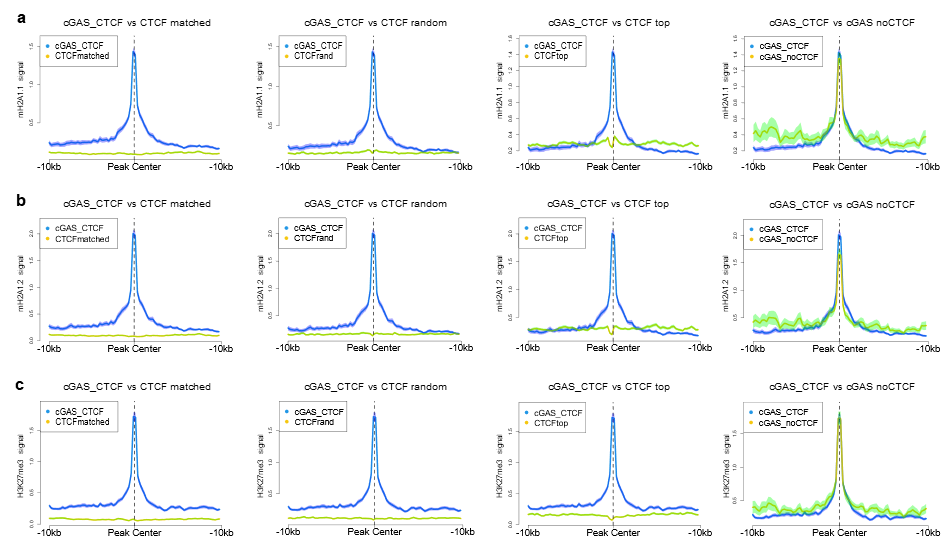

Fig. S6.

Average ChIP signal enrichment profiles for mH2A1.1 (a), mH2A1.2 (b), and H3K27me3 (c) across cGAS peaks bound by CTCF (blue lines in all plots). Green lines correspond to ChIP signals across CTCF peaks, non-overlapping cGAS peaks (separated into three enrichment tiers: matched, random, and top; columns 1–3, respectively), or cGAS peaks not bound by CTCF (column 4). Shaded areas represent standard error across each ChIP signal bin (250 bp).

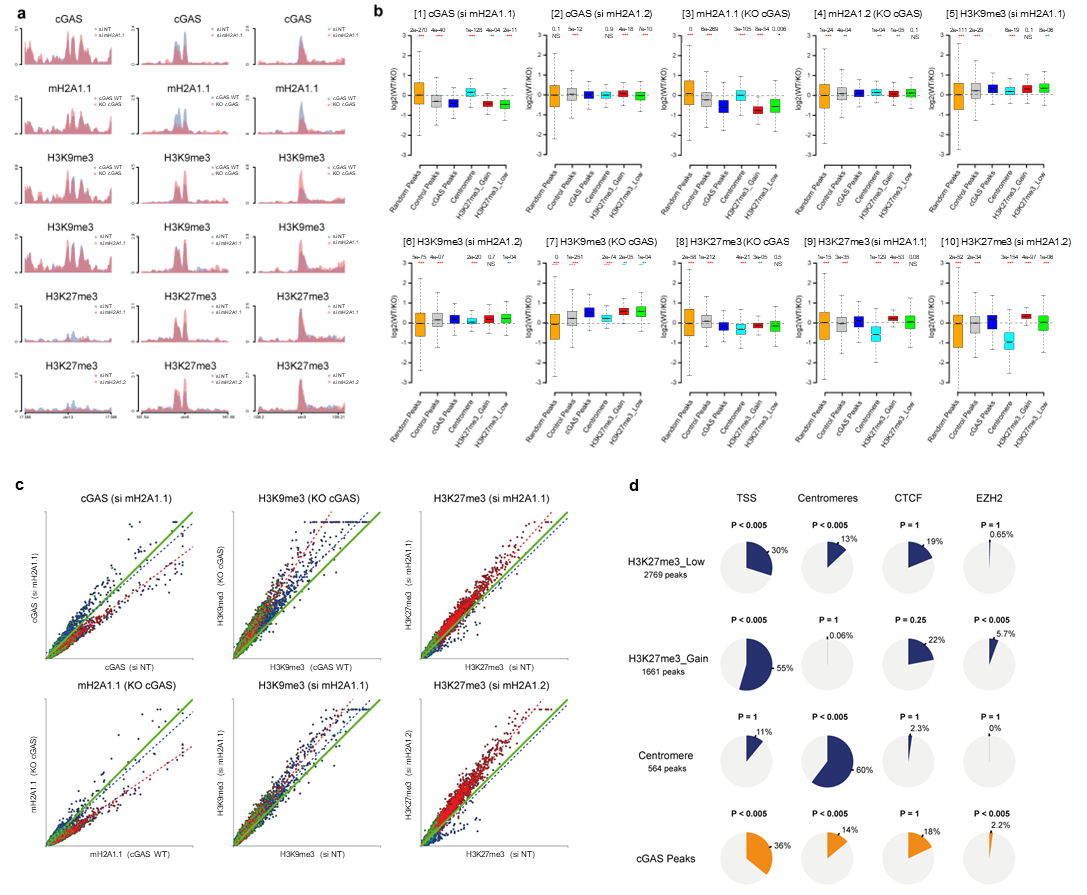
Fig. S7.

a, Representative examples of cGAS peaks illustrating variation in ChIP-seq enrichment levels in WT (blue) and KO/siRNA (red) conditions. Each vertical panel shows cGAS, mH2A1.1, mH2A1.2, H3K27me3, and H3K9me3 enrichment profiles over a 10 kb genomic region centered on the peak summit. Clusters are ordered horizontally and correspond to centromeric-associated (left), H3K27me3-associated (middle), and H3K9me3-associated (right) behaviors (Fig. 6c,d). b, Boxplots showing ChIP-seq enrichment fold change (log₂(WT/KO)) across perturbations. Panel titles indicate targeted ChIP-seq levels (WT) and perturbations (KO or siRNA) in parentheses. ChIP signal levels represent average enrichment over a 500 bp region centered on peak summits. P-values were derived by comparing distributions of each cluster to all cGAS peaks (blue) using a Wilcoxon two-sided test. c, Scatterplots depicting major variation patterns in cGAS binding regions upon: cGAS depletion (impact on H3K9me3 [top mid] and mH2A1.1 [bottom left]), mH2A1.1 depletion (impact on cGAS [top left], H3K27me3 [top right], and H3K9me3 [bottom mid]), mH2A1.2 depletion (impact on H3K27me3 [bottom right]). Data are color-coded: centromeres (blue), H3K27me3 (red), H3K9me3 (green). ChIP signal levels reflect average enrichment over 500 bp regions centered on peak summits, with WT values on the x-axis. Dotted lines represent linear models fitted to centromeric-associated (blue) or H3K27me3-associated (red) cGAS peaks. d, Pie charts showing the proportion of cGAS peaks in each cluster overlapping annotated TSSs, centromeres, CTCF peaks, or EZH2 peaks. P-values were calculated using a permutation test (200 permutations) against randomized genomic coordinates, with overlap thresholds set to 1 kb.

Table S1.

MS/MS analysis of cGAS-Flag-HA tandem affinity chromatography.

**References**

61.	Posavec Marjanović, M. *et al.* MacroH2A1.1 regulates mitochondrial respiration by limiting nuclear NAD+ consumption. *Nat. Struct. Mol. Biol.* **24**, 902–910 (2017).

62.	Tanaka, Y. *et al.* Expression and purification of recombinant human histones. *Methods* **33**, 3–11 (2004).

63	Tachiwana, H. *et al.* Structural basis of instability of the nucleosome containing a testis-specific histone variant, human H3T. *Proc. Natl. Acad. Sci. U. S. A.* **107**, 10454–10459 (2010).

64.	Punjani, A., Rubinstein, J. L., Fleet, D. J. & Brubaker, M. A. cryoSPARC: algorithms for rapid unsupervised cryo-EM structure determination. *Nat. Methods* **14**, 290–296 (2017).

65.	Thorne, R. E. Hypothesis for a mechanism of beam-induced motion in cryo-electron microscopy. *IUCrJ* **7**, 416–421 (2020).

66.	Ma, C.-W., Zhang, J., Dong, X.-Q. & Lu, J.-X. Amyloid structure of high-order assembly of Leucine-rich amelogenin revealed by solid-state NMR. *J. Struct. Biol.* **206**, 29–35 (2019).

67.	Sanchez-Garcia, R. *et al.* DeepEMhancer: a deep learning solution for cryo-EM volume post-processing. *Commun. Biol.* **4**, 874 (2021).

68.	Adams, P. D. *et al.* PHENIX: a comprehensive Python-based system for macromolecular structure solution. *Acta Crystallogr. D Biol. Crystallogr.* **66**, 213–221 (2010).

69.	Emsley, P., Lohkamp, B., Scott, W. G. & Cowtan, K. Features and development of Coot. *Acta Crystallogr. D Biol. Crystallogr.* **66**, 486–501 (2010).

70.	Alexander, N., Woetzel, N. & Meiler, J. bcl::Cluster : A method for clustering biological molecules coupled with visualization in the Pymol Molecular Graphics System. *IEEE Int. Conf. Comput. Adv. Bio Med. Sci. Proc. IEEE Int. Conf. Comput. Adv. Bio Med. Sci.* **2011**, 13–18 (2011).

71.	Goddard, T. D. *et al.* UCSF ChimeraX: Meeting modern challenges in visualization and analysis. *Protein Sci. Publ. Protein Soc.* **27**, 14–25 (2018).

72.	Recoules, L. *et al.* The histone variant macroH2A1.1 regulates RNA polymerase II-paused genes within defined chromatin interaction landscapes. *J. Cell Sci.* **135**, jcs259456 (2022).

73.	Dardenne, E. *et al.* Splicing switch of an epigenetic regulator by RNA helicases promotes tumor-cell invasiveness. *Nat. Struct. Mol. Biol.* **19**, 1139–1146 (2012).

74.	Machida, S. *et al.* Exploring histone loading on HIV DNA reveals a dynamic nucleosome positioning between unintegrated and integrated viral genome. *Proc. Natl. Acad. Sci. U. S. A.* **117**, 6822–6830 (2020).

75.	Molina, O. *et al.* Epigenetic engineering reveals a balance between histone modifications and transcription in kinetochore maintenance. *Nat. Commun.* **7**, 13334 (2016).

76.	Luo, Y. *et al.* New developments on the Encyclopedia of DNA Elements (ENCODE) data portal. *Nucleic Acids Res.* **48**, D882–D889 (2020).

77.	Reiff, S. B. *et al.* Author Correction: The 4D Nucleome Data Portal as a resource for searching and visualizing curated nucleomics data. *Nat. Commun.* **13**, 6561 (2022).
